# TMEM119+ microglia MHC class I restricted antigen presentation impacts CD8 T cell memory, effector status, and blood-brain barrier disruption during neurotropic virus infection

**DOI:** 10.64898/2026.05.08.722741

**Authors:** Marina Seady, Mark A. Maynes, Javonte S. Thelwell, Fang Jin, Michael J. Hansen, Hadley E. Jensen, Ryann K. Witter, Carley A. Owens, Asma Hassani, Cody L. Lewis, Michael D. Forston, Aaron J. Johnson

## Abstract

The impact of microglia antigen presentation on CNS infiltrating CD8 T cells responses during neurotropic virus infection has been difficult to define. Using Theiler’s murine encephalomyelitis virus (TMEV) infection of neurons as a model system, our laboratory has previously determined that H-2D^b^ restricted, but not H-2K^b^ restricted CD8 T cells are required for viral clearance, demonstrating the role of discrete MHC class I alleles. To determine the extent microglia antigen presentation impacts brain-infiltrating CD8 T cells, our laboratory generated novel single MHC class I conditional knockout mice in which H-2K^b^ or H-2D^b^ can be specifically deactivated in TMEM119+ microglia with tamoxifen administration. During TMEV infection, conditional knockout of H-2K^b^ in microglia reduced antigen-specific CD8 T cell proliferation in the brain. Meanwhile, mice with deletion of D^b^ in microglia had reduced levels of perforin in antigen-specific CD8 T cells. Furthermore, microglia specific deletion of H-2D^b^ reduced CD8 T cell numbers in the brain and preserved blood-brain barrier (BBB) integrity. Microglial D^b^ restricted antigen presentation was also essential for the reactivation of CD8 tissue resident memory (TRM) cells and BBB integrity during memory recall responses. These findings further our understanding of how brain infiltrating antiviral CD8 T cell responses are impacted by microglia, as well as define how this cellular interaction contributes to BBB disruption during neuroinflammation. These findings also have high significance to our understanding of how microglia impact CD8 TRM cell populations that reside in the brain long after virus infection is cleared.

**Graphical abstract:** 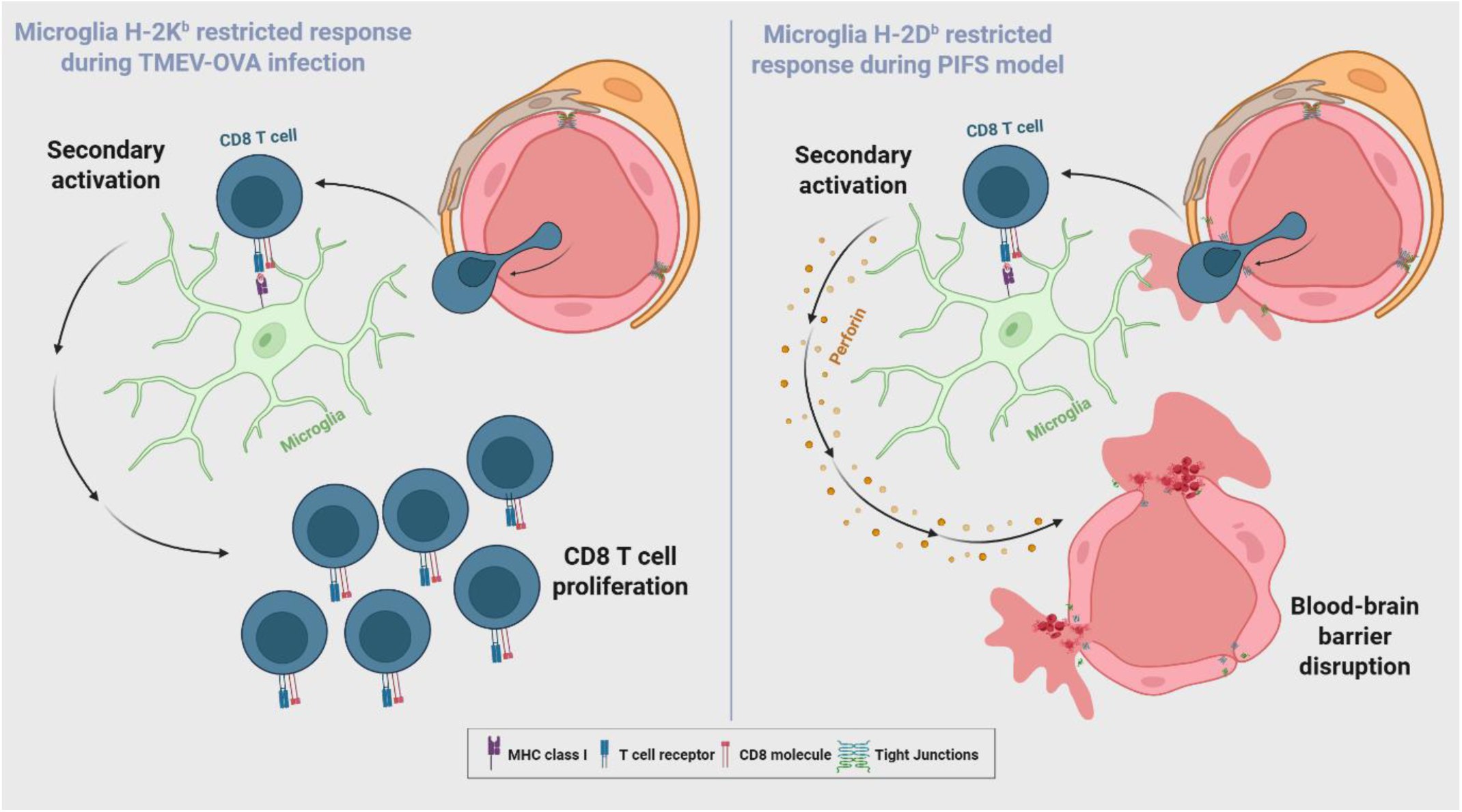

## Introduction

Neurotropic viral infections are responsible for acute and long-lasting neurological manifestations, resulting in burdens on healthcare systems worldwide ^1, 2^. The contribution of long-lived memory T cells in the CNS well after viruses are cleared also contribute to chronic neuroinflammation through means poorly understood ^3, 4, 5, 6, 7^. Central to chronic neuroinflammation is microglial activation, which in turn modifies T cell responses against neurotropic viruses ^8, 9^. Microglia have the capacity to influence neurotropic virus infections through the upregulation of phagocytosis machinery, enhanced expression of proinflammatory molecules, and uptake of cellular debris ^10, 11^. However, the role of microglia activation is not limited to innate immune function due to their increased expression of antigen processing and presentation molecules ^8, 12, 13, 14, 15^. As such, it is assumed microglia become fully competent antigen presenting cells which boost antiviral T cell responses in the brain ^8, 12, 13, 15^. However, it remains unclear how T cell responses are altered acutely and long term when microglia are not directly infected with virus ^16^.

Theiler’s murine encephalomyelitis virus (TMEV) provides a neurotropic virus infection model in which the contribution of discrete MHC class I molecules are well defined ^17, 18,.19^. H-2D^b^ restricted, but not H-2K^b^ restricted CD8 T cells are required for viral clearance, demonstrating the role of discrete MHC class I alleles ^17^. To define how myeloid cells impact T cell responses against neurotropic virus infection through H-2K^b^ and H-2D^b^, Goddery *et al*. evaluated the impact of conditionally deleting discrete MHC class I molecules in CX3CR1+ cells following Theiler’s murine encephalomyelitis virus (TMEV) infection ^8^. This conditional silencing of MHC class I restricted antigen presentation resulted in a dampened CD8 T cell response and reduced brain atrophy post TMEV infection ^8^. In agreement with these findings, Moseman *et al.* analyzed vesicular stomatitis virus (VSV) infection using a tracer for olfactory sensory neurons (OSNs) in the olfactory bulb ^15^. 6 days post VSV infection, fragments of labeled OSNs were found within CX3CR1+ cells, also demonstrating in principle that microglia could acquire and present antigen from virus-infected neurons ^15^.

In both of these studies, myeloid cells were determined to impact CD8 T cell responses, potentially through the cross-presentation of virus antigen. However, it has remained difficult to define the extent microglia enhance CD8 T cell responses in the brain through MHC class I restricted antigen presentation due to the lack of microglia-specific tools. Approaches using CX3CR1 Cre mice and cell depletion strategies targeting the colony-stimulating factor 1 receptor (CSF1R) have been informative, but perivascular macrophages and infiltrating myeloid cells are common off target effects of myeloid cell ablation strategies that need consideration when interpreting results ^20, 21, 22^.

Using a novel single MHC class I Cre Lox approach developed by our laboratory in which H-2D^b^ and H-2K^b^ can be deactivated in TMEM119+ cells, we sought to understand the impact of microglia MHC class I restricted antigen presentation on acute antiviral CD8 T cell responses and long lived CD8 T resident memory cells post virus clearance. Using TMEV and recombinant TMEV encoding the model OVA (TMEV-OVA) antigen, we were able to analyze virus antigen-specific CD8 T cells restricted to D^b^ class I and K^b^ class I molecules, respectively. Our data supports that K^b^ and D^b^ have discrete roles in modifying brain infiltrating CD8 T cells, and D^b^ restricted antigen presentation on microglia is essential for CD8 tissue resident memory (TRM) recall. We also put forward evidence that microglia antigen presentation promotes CD8 T cell induced BBB disruption.

## Methods

### Animals

Male and female C57BL/6J wildtype (WT) (stock no. 000664) and Tmem119-creER^T^^2^ mice (stock no. 031820) were purchased from The Jackson Laboratory. H-2K^b^ LoxP and H-2D^b^ LoxP mice were generated by our program, as previously described, in the C57BL/6 background. H-2K^b^ LoxP and H-2D^b^ LoxP mice were crossed with Tmem119-Cre^ERT^^2^ mice to generate Tmem119 K^b^ cKO and Tmem119 D^b^ cKO mice that express Cre under the Tmem119 promoter and undergo microglia-specific targeted gene deletion of H-2K^b^ and H-2D^b^ molecules. Mice were bred and maintained under specific pathogen-free conditions at the Mayo Clinic Animal Facility. In the studies, 10–15-week-old mice were used for all experiments with appropriate age- and sex-matched controls. All experiments were conducted in accordance with guidelines from the National Institutes of Health and with the approval of the Mayo Clinic Institutional Animal Care and Use Committee.

### Tamoxifen administration

Tamoxifen (Sigma-Aldrich, St. Louis, MO) was administered in corn oil (Sigma-Aldrich, St. Louis, MO) at a concentration of 20mg/mL. Tamoxifen was dissolved in corn oil by constant agitation overnight at 37°C in the dark. Tmem119 K^b^ cKO mice were administered 75 mg/kg tamoxifen intraperitoneally at 8 weeks of age once a day for five consecutive days using a 26-gauge 3/8” beveled needle. Tmem119 D^b^ cKO were administered 75 mg/kg tamoxifen intraperitoneally at 8 weeks of age for 5 consecutive days followed by one week of recovery without injections, and additional 5 days of tamoxifen injections daily. The differential tamoxifen regime was defined after testing for the expression of MHC class I molecules on TMEM119+ microglia after 1 week of injections (sufficient for Tmem K^b^ cKO mice) or 2 weeks of injections (sufficient for Tmem119 D^b^ cKO mice). Post tamoxifen recovery times of 1 week was incorporated prior to experimental use of mice in experiments.

### TMEV and TMEV-OVA infection

The Daniel’s strain of Theiler’s murine encephalomyelitis (TMEV) was prepared as previously described ^19^. TMEV-XhoI-OVA8 [TMEV-ovalbumin (OVA)] was generated and prepared by our group, as previously described ^23^. At 10–12 weeks of age, mice were anaesthetized with 1–2% isoflurane and infected intracranially with 2 × 10^6^ PFU (10 µl) of the Daniel’s strain of TMEV or 2 × 10^5^ PFU of TMEV-OVA. Virus was delivered to the right hemisphere of the brain via automatic 1 mL Hamilton syringe (Hamilton). Mice were euthanized for high-dimensional flow cytometry of tissues at 7 days post-infection (dpi) or according to experimental design displayed in every figure.

### FTY720 treatment and Bromodeoxyuridine (BrdU) administration

FTY720 (Sigma, SML0700–5mg) was diluted to 0.15mg/mL in water such that a 100μL intraperitoneal injection would be the equivalent of a 0.5mg/kg dose assuming a 30g mouse. FTY720 was administered in this way twice a day over the course of appropriate experiments to randomized, age matched Tmem119 K^b^ cKO and Cre negative littermate controls. 100μL of 10mg/mL BrdU (BD Pharmingen, Cat #51–2420KC) solution was administered intraperitoneally 5 days post intracranial TMEV-OVA infection. For BrdU staining, manufacturer’s instructions were followed according to the kit.

### Intravascular labeling

For Intravascular labeling experiments, 200μL total volume (3μg of FITC αCD45 (Cat #FITC 35–0451-U100, Tonbo) diluted in PBS) was injected into the tail vein 24 hours before euthanasia.

### FITC-albumin leakage assay

Mice were intravenously injected with 100 µL of a solution containing 66 kDa FITC-conjugated albumin diluted in PBS 1 h prior to euthanasia to visualize regions of blood–brain barrier disruption and vascular leakage. Mice were not perfused before euthanasia. Brains were paraformaldehyde-fixed for 48 h following mouse euthanasia. FITC-albumin injected brains were embedded in 4% agarose and sliced with a vibratome. All brain slices were mounted on microscope slides and stored at 4°C. Images were acquired using the Echo Revolution microscope. Analysis was performed using whole-slice thresholding followed by thresholding of the FITC channel acquired by ImageJ (National Institutes of Health). Results are shown as % of FITC per total area.

### Reactivation of virus antigen-specific CD8 T cells through peptide injection

Between 55-65 days post-TMEV infection, mice were intravenously injected with 0.1 mg (from a solution of 1mg/ml in PBS) of VP2_121–130_ (FHAGSLLVFM) or control E7 (RAHYNIVTF) peptide (GenScript, Piscataway, NJ, USA) into the tail vein. Both of these peptides are MHC I (D^b^) restricted.

### Single-cell suspension

Mice were euthanized via isoflurane per institutional animal care and use committee (IACUC) guidelines and transcardially perfused with ice-cold PBS. Perfused brains were processed following a previously published protocol designed for enrichment of brain infiltrating leukocytes using the Dounce homogenization method followed by 30–38% Percoll (Cat# P1644–1L, Sigma, St. Louis, MO) gradient centrifugation. Thymus and spleen samples were processed using mechanical disruption with the rough side of two microscope slides through a 70 µM filter. Blood was collected (100 µL) into 1 ml heparin (100 U/mL). Spleens underwent ammonium-chloride-potassium (ACK) RBC lysis for 3 min and blood samples for 1 min. All samples were filtered using 70 µM filters, and resulting cells were counted on the hemocytometer (Hausser Scientific, Horsham, PA) using trypan blue to exclude dead cells (Gibco, Gaithersburg, MD). Alive cells were then stained and ran on a flow cytometer.

### Flow cytometry

When applicable, samples were stained with 50 µL of a 1:50 dilution of D^b^: VP2_121-130_ APC-labeled tetramer or a 1:50 dilution of K^b^: OVA_257-264_ APC-labeled tetramer (NIH Tetramer Core Facility, Emory University). For tetramer staining, we incubated cells with only the tetramer for 30 minutes at room temp before proceeding with staining with viability dye and the rest of antibodies. Zombie UV dye (Cat# 423108, Biolegend) was used at a 1:1000 concentration to identify and exclude non-viable cells. The Zombie UV dye was stained separately for 20 minutes at room temp and washed off before staining with the rest of the master mix. The following antibodies were used at a 1:50 dilution: PE-Cy7 Tmem119 (ThermoFisher, Cat# 25-6119-82), BV510 CD4 (BioLegend, Cat# 100559), PE H-2K^b^ (BD Pharmingen, Cat# 5535570), PE H-2D^b^ (Biolegend, Cat# 111508), BV650 IFNγ (Biolegend, Cat# 505831), FITC perforin (Biolegend, Cat# 154310) and Alexa Fluor 700 TNFα (Biolegend, Cat# 506338).The following antibodies were used at a 1:100 dilution: BUV615 NK-1.1 (BD Biosciences, Cat# 751111), BUV395 F480 (BD Horizon, Cat# 565614), BUV737 TCRβ (BD OptiBuild, Cat# 612821), BV421 CXCR3 (Biolegend, Cat# 1265522), BV570 CD44 (Biolegend, Cat# 103037), BV605 CD11C (Biolegend, Cat# 117333), PerCP Ly6C (Biolegend, Cat# 128027), APC FIRE 750 CD62L (Biolegend, Cat# 104450), APC FIRE 810 CD19 (Biolegend, Cat# 115578), BV421 CD103 (Biolegend, Cat#121422) and Pacific Blue CX3CR1 (Biolegend, Cat#149037). The following antibodies were used at a 1:200 dilution: BV711 Ly-6G (BioLegend, Cat# 127643), Spark Blue 550 CD8a (BioLegend, Cat# 100780), BUV805 MHC II (BD OptiBuild, Cat# 748844), BB700 CD24 (BD Biosciences, Cat# 746122) and Spark NIR 685 CD69 (Biolegend, Cat# 104558). The following antibodies were used at a 1:1000 dilution: PE-CF594 CD45 (BioLegend, Cat# 562420) and PE-Cy5 CD11b (TONBO Biosciences, Cat# 55-0112-U100). To all master mixes of antibodies, FC block (Cat# 553141, BD) was added at a 1:100 dilution to prevent non-specific binding. The single-cell suspensions were next fixed and permeabilized with 250 μL of Cytofix/Cytoperm fixation buffer (BD Biosciences, Franklin Lakes, NJ) for 20 min at 4 °C. Antibodies that tag intracellular proteins were used in a separate master mix, and cells were stained after permeabilization. Across all staining conditions, cells were analyzed on the spectral high parameter flow cytometer Cytek Aurora (Cytek, Fremont, CA) and data was analyzed on FlowJo 10.1 (FlowJo LLC, Ashland, OR). High dimensional analysis including UMAP analysis was also performed using FlowJo software using DownSample and UMAP_R plugins. For BrdU staining, manufacturer’s protocol was followed.

### Parabiosis

Two female age-matched mice of similar weight were selected. Mice were either littermates or co-housed for a minimum of 2 weeks prior to surgery. Anesthesia was induced and maintained with ketamine/xylazine (100/10 mg/kg; maintenance dose 30/3 mg/kg). Ophthalmic ointment was placed on eyes. Mice were tested for reflex responses to ensure anesthetic depth. For analgesia during surgery, we administered intraperitoneal carprofen and buprenorphine-SR subcutaneously (5 mg/kg and 1 mg/kg, respectively). Animals were shaved on the flank and prepped for surgery with Betadine swabs and sterile drapes. Using sharp scissors, a superficial epidermal incision was continuously made along the flank starting at 0.5 cm rostral of olecranon to 0.5 cm superior to patella. Skin absorbable 3–0 silk sutures were used to join elbow and knee joints (Ethicon Inc.). Simple continuous suture was performed with absorbable 4–0 vicryl suture (Ethicon Inc.) for closer of incision. Postoperatively, we administered 1 ml of 0.9% NaCl fluids subcutaneously and analgesic carprofen intraperitoneally and buprenorphine-SR subcutaneously. Mice were placed on heating pads and observed for a minimum of 3 h after surgery. Prophylactically, mice were treated with antibiotic Enrofloxacin (50 mg/kg) 10 days postoperatively. Parabiont pairs underwent a minimum of 3 weeks of healing and acclimation time before experimental use. The procedure was performed according to the IACUC guidelines at Mayo Clinic.

### Magnetic Resonance imaging (MRI)

MRI images were acquired and processed as previously described ^24, 25^. A Bruker Avance II 300-MHZ (7T) vertical-bore small-animal animal system was used to conduct T1-weighted scans (Bruker Biospin, Billerica, MA, USA) following an intraperitoneal injection of gadolinium 15 minutes before the MRI. Anesthesia was induced with isoflurane and maintained for the duration of the scan. Respiratory rate and temperature were monitored. Analysis and quantification of the gadolinium leakage into the brain was performed using Analyze 12.0 software (Biomedical Imaging Resource, Mayo Clinic). The analysis of gadolinium leakage in the T1-weight MRIs was performed blinded.

### Statistical analysis

When comparing results from two groups, an unpaired Student’s t-test was used; when comparing results from two groups of paired mice, a paired Student’s t-test was used; and when comparing results from three groups, a one-way ANOVA using Tukey’s post hoc multiple comparisons test was used. A two-way ANOVA was used when comparing statistically significant differences between the means of three or more independent groups that had been split on two variables, using Šidák’s test for multiple comparisons. All measurements in all experiments were taken from distinct samples. Normal distribution was tested using the Shapiro–Wilk test. All parabiont data employed pairwise comparisons. All statistical analyses were performed using Prism 10.0 (GraphPad Software), Excel or R. Data are presented as the mean ± standard deviation (SD), and all P-values are shown. Non-significant P-values (P > 0.05) are marked.

## Results

### Establishment of microglia H-2K^b^ and H-2D^b^ conditional knockout (cKO) mice

The challenge of dissecting microglial contributions to neurologic disease stems from the use of strategies that also target additional myeloid cells, namely perivascular macrophages, infiltrating monocytes, dendritic cells, macrophages and other circulating myeloid cells ^20, 22, 26^. Tmem119 expression is highly specific to microglia, enabling targeted deletion employing Cre-Lox technologies in mouse models ^27, 28^. Our laboratory has developed tamoxifen-inducible conditional knockout strategies to dissect the contribution of two discrete MHC class I molecules, H-2K^b^ and H-2D^b^, on microglia. The Tmem119 K^b^ cKO mice is designed to have H-2K^b^ ablation in microglia. Likewise, Tmem119 D^b^ cKO mice is designed to have targeted inactivation of microglia H-2D^b^ expression. Throughout this study, these mice with the ablation of K^b^ or D^b^ on TMEM119+ microglia will be compared to K^b^ Cre- littermate control or D^b^ Cre- littermate controls, respectively. To confirm H-2K^b^ and H-2D^b^ expression was specifically ablated in TMEM119+ microglia, we administered tamoxifen and then intracranially infected mice with TMEV. Microglial expression of H-2K^b^ and H-2D^b^ in TMEM119+ cells in brains isolated from TMEV infected animals was then assessed using spectral flow cytometry. As intended, we observed a marked reduction of H-2K^b^ or H-2D^b^ in TMEM119+ cells in these animals (Figure 1 A, C). To further validate specificity of MHC class I knockdown, we assessed H-2K^b^ (Figure 1B) and H-2D^b^ (Figure 1D) expression in blood immune cell populations and confirmed there was no difference between Cre- and Cre+ mice in both K^b^ and D^b^ cKO mice.

**Figure 1.**
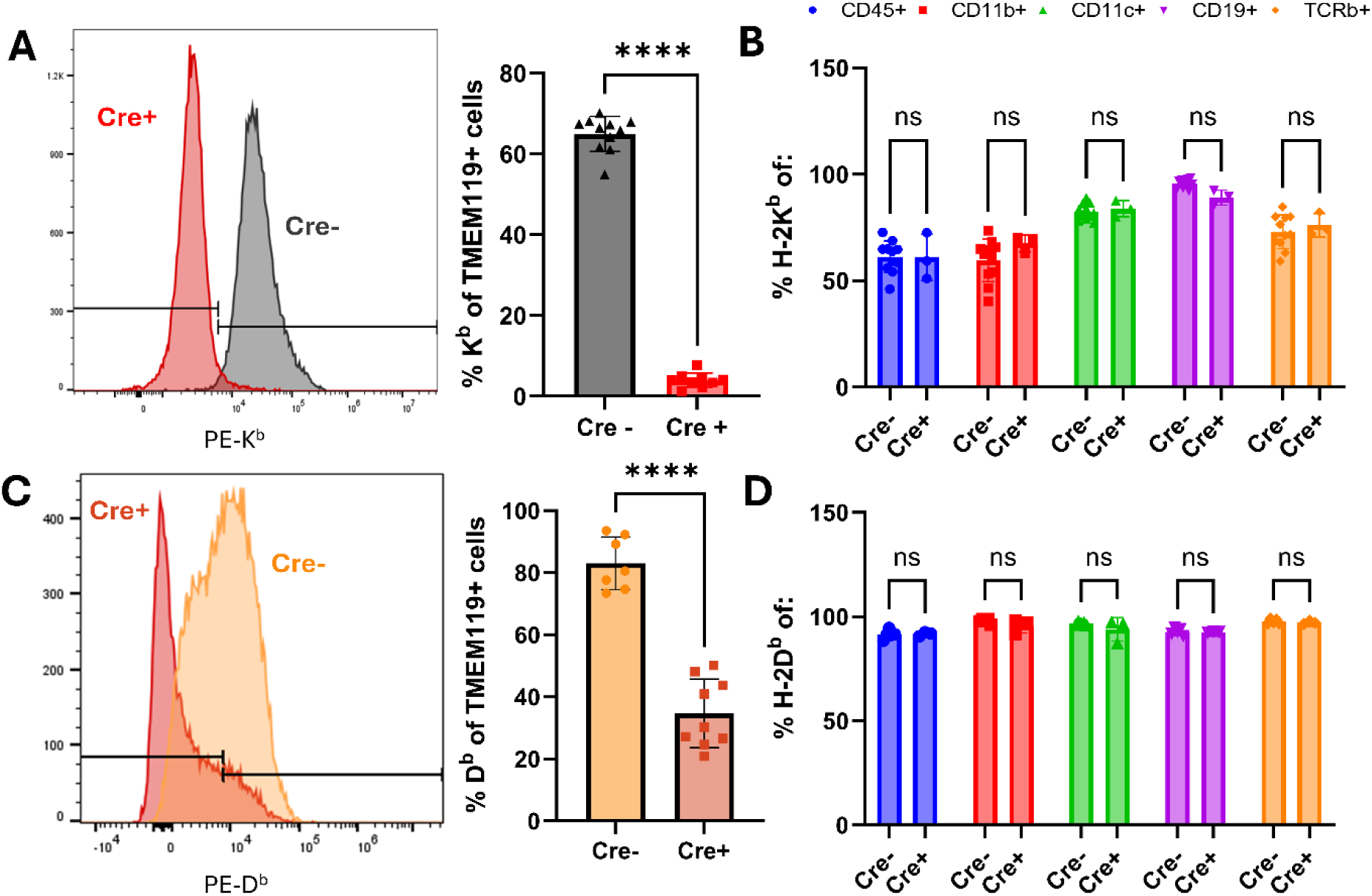
Development of TMEM119+ microglia MHC class I molecule conditional knockout mice. (A) Representative histograms and percentages of K^b^ expression of TMEM119+ cells in the brain in Tmem119 K^b^ cKO mice compared to Cre- littermate controls 7 days post TMEV infection, followed by (B) H-2K^b^ expression in several immune cell populations in the blood. (C) Representative histograms and percentages of D^b^ expression of TMEM119+ cells in the brain in Tmem119 D^b^ cKO mice compared to Cre-littermate controls 7 days post TMEV infection, followed by (D) H-2D^b^ expression in several immune cell populations in the blood. Designation of symbols is as follows: ns or not shown for p > 0.05, * for p ≤ 0.05, ** for p ≤ 0.01, *** for p ≤ 0.001, **** for p ≤ 0.0001. Data presented as mean +/- SD.

We next wanted to determine if conditional deletion of K^b^ and D^b^ MHC class I molecules on TMEM119+ cells impacted CD8 T cell development and repertoire diversity. We analyzed TCR Vβ usage in the spleens of naïve animals for Tmem119 K^b^ cKO (Figure 2A) and Tmem119 D^b^ cKO mice (Figure 2B). We did not observe differences between Tmem119 K^b^ and D^b^ cKO mice and Cre- littermate controls. As a measurement of thymocyte output, we analyzed the percentages of single positive CD8 and CD4 T cells for both cKOs (Figure 2C and 2D). We did not observe differences between groups, demonstrating T cell development in the thymus is not affected by targeted ablation of MHC class I molecules in microglia.

**Figure 2.**
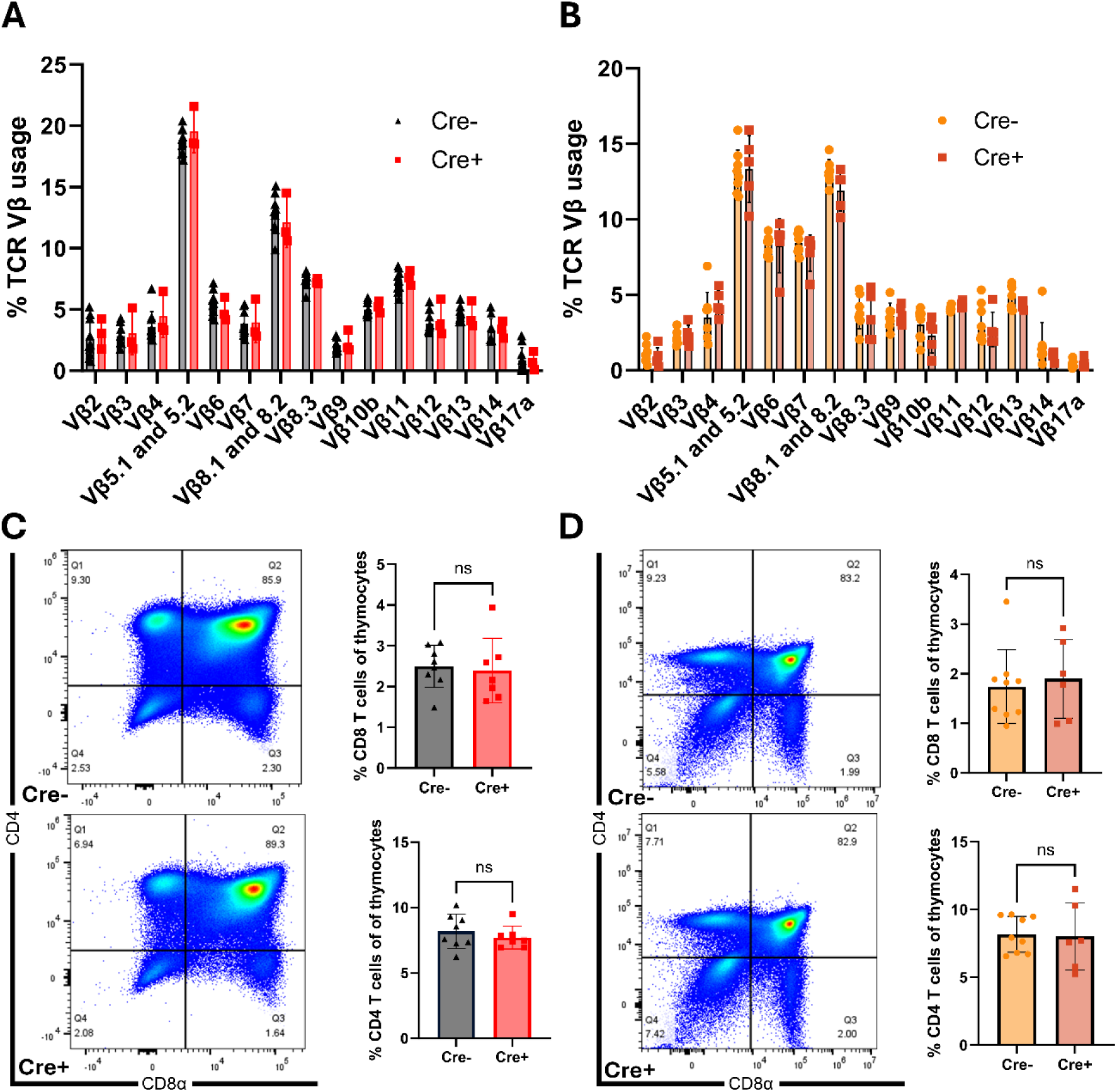
Normal T-cell development in naive Tmem119 K^b^ cKO (gray and pink) and Tmem119 D^b^ cKO mice (orange and red) in comparison to Cre negative littermates. (A, B) Splenic TCR repertoire is not affected by cell-specific H-2K^b^ and H-2D^b^ deletion. Splenocytes were isolated from naïve 10-week-old animals (N = 3-7 per group) and stained with anti-CD45, anti-TCRβ, anti-CD4, and anti-CD8 antibodies, as well as antibodies specific to each of the T-cell receptor Vβ regions listed. (C, D) Representative plots and quantification show no disruption of thymocyte development in Tmem119 K^b^ cKO mice and Tmem119 D^b^ cKO in comparison to Cre negative littermates. Designation of symbols is as follows: ns or not shown for p > 0.05. Data presented as mean +/- SD.

### Conditional deletion of H-2K^b^ on microglia reduces antigen specific CD8 T cell expansion in the brain 7 days post TMEV-OVA infection

After validating our conditional knockout approach, we proceeded to investigate the extent microglia H-2K^b^ was impacting CD8 T cell responses in the brain. Using our targeted Cre-Lox approach that enables conditional deletion of H-2K^b^ in TMEM119+ cells (Figure 3A), we infected mice with recombinant TMEV-OVA virus which encodes the model antigen SIINFEKL presented in the context of the K^b^ class I molecule ^23^. We evaluated the impact on the ensuing brain infiltrating immune response at 7 days post infection (Figure 3B). We observed a marked reduction in the frequencies of several immune populations such as NK cells, dendritic cells, macrophages and inflammatory monocytes (Figure 3C and Figure S1). Of major note, K^b^:OVA epitope specific CD8 T cells were markedly reduced in Tmem119 K^b^ cKO mouse brains post infection, indicating that microglia K^b^ restricted antigen presentation was required for complete CD8 T cell expansion.

**Figure 3.**
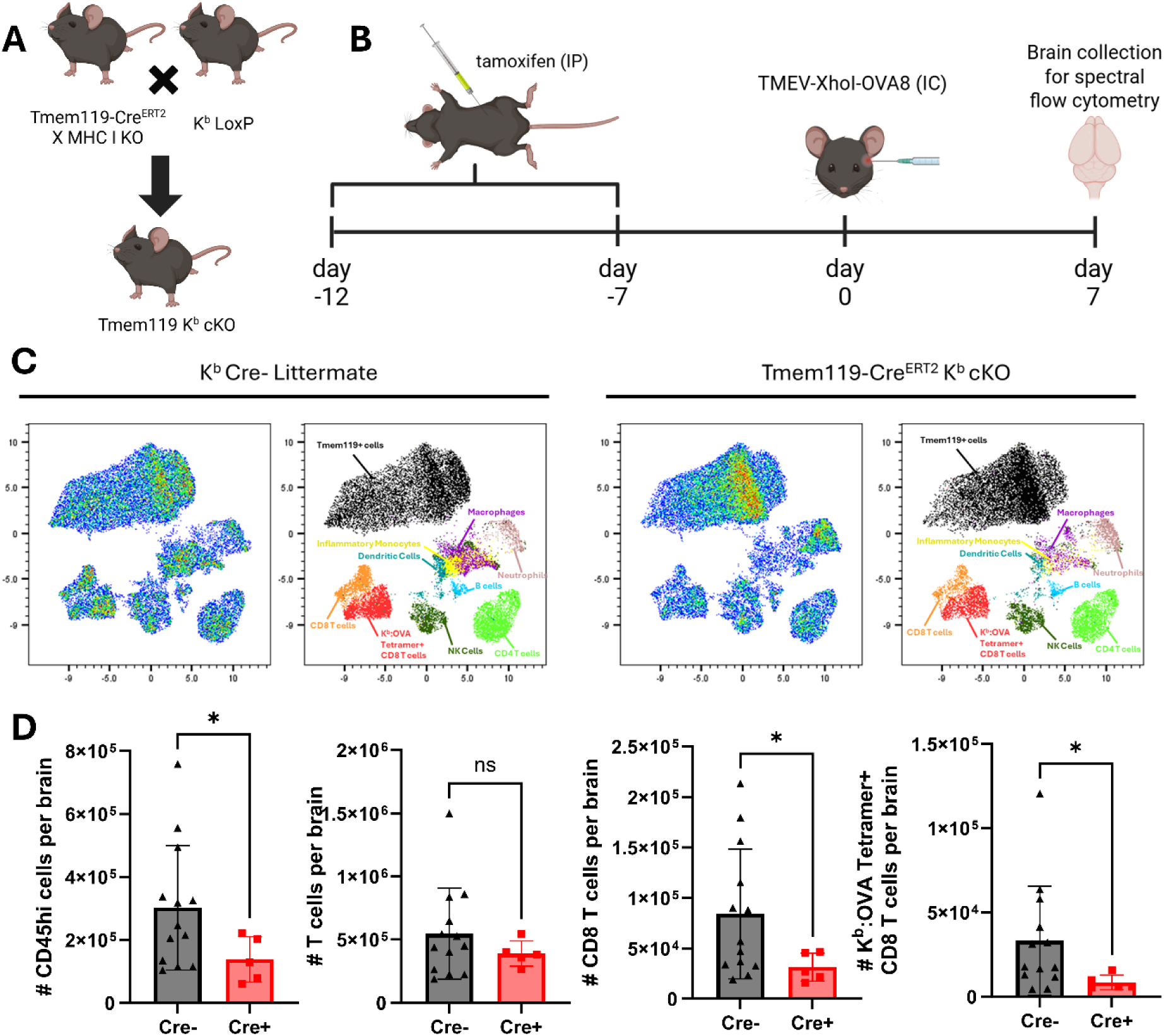
Tmem119 K^b^ cKO mice exhibit decreased K^b^:OVA epitope specific CD8 T cell numbers at brain 7 days post TMEV-OVA infection. (A) Schematic depicting the Cre-LoxP strategy. Tmem119-Cre^ERT^^2^ mice were crossed to transgenic H-2K^b^ LoxP mice. (B) Experimental design showing mice were treated with tamoxifen to activate cre for 5 days, once a day. Mice were infected with TMEV-OVA and euthanized 7 days post infection. Brains were analyzed using spectral flow cytometry. (C) Representative uniform manifold approximation and projection (UMAP) key, colored to highlight the different cell subsets in the brain for both K^b^ Cre- Littermate and Tmem119-CreERT2 K^b^ cKO mice. (D) Absolute counts of CD45hi, T cells, CD8 T cells and K^b^:OVA Tetramer+ CD8 T cells per brain. Designation of symbols is as follows: ns or not shown for p > 0.05, * for p ≤ 0.05, ** for p ≤ 0.01, *** for p ≤ 0.001, **** for p ≤ 0.0001. Data presented as mean +/-SD.

### CD8 T cell proliferation is attenuated when K^b^ is conditionally ablated in microglia during acute infection

To define the mechanism by which H-2K^b^ expressed in microglia enhances CD8 T cell responses in the brain, we evaluated the extent cell proliferation was a factor. We pulsed mice with intraperitoneal administered BrdU to evaluate cell proliferation. BrdU is incorporated into newly synthesized DNA by cells entering and progressing through the S (DNA synthesis) phase of the cell cycle, a process detectable by anti-BrdU antibodies using spectral flow cytometry. Substantial CD8 T cell infiltration of the brain occurs at 7 days post TMEV (and TMEV-OVA) infection. However, virus antigen-specific CD8 T cells first infiltrate the brain at day 4 post infection ^29^. We therefore designed our experiment to enable this first wave of CD8 T cells at day 4 to enter the brain (Figure 4A). We next treated mice with FTY720 to sequester T cells within the secondary lymphoid organs and prevent further brain infiltration of antiviral CD8 T cells. FTY720 treatment therefore allowed us to analyze T cell proliferation specifically in the brain. Following this experimental design, we confirmed that FTY720 treatment resulted in decreased T cell numbers in the blood (Figure S2). Importantly, we determined that the percentage of BrdU+ T cells and antiviral K^b^:OVA antigen specific CD8 T cells were significantly decreased in the brains of Tmem119 K^b^ cKO mice (Figure 4C). This confirmed microglia K^b^ class I restricted antigen presentation was required for optimal CD8 T cell proliferation in the brain.

**Figure 4.**
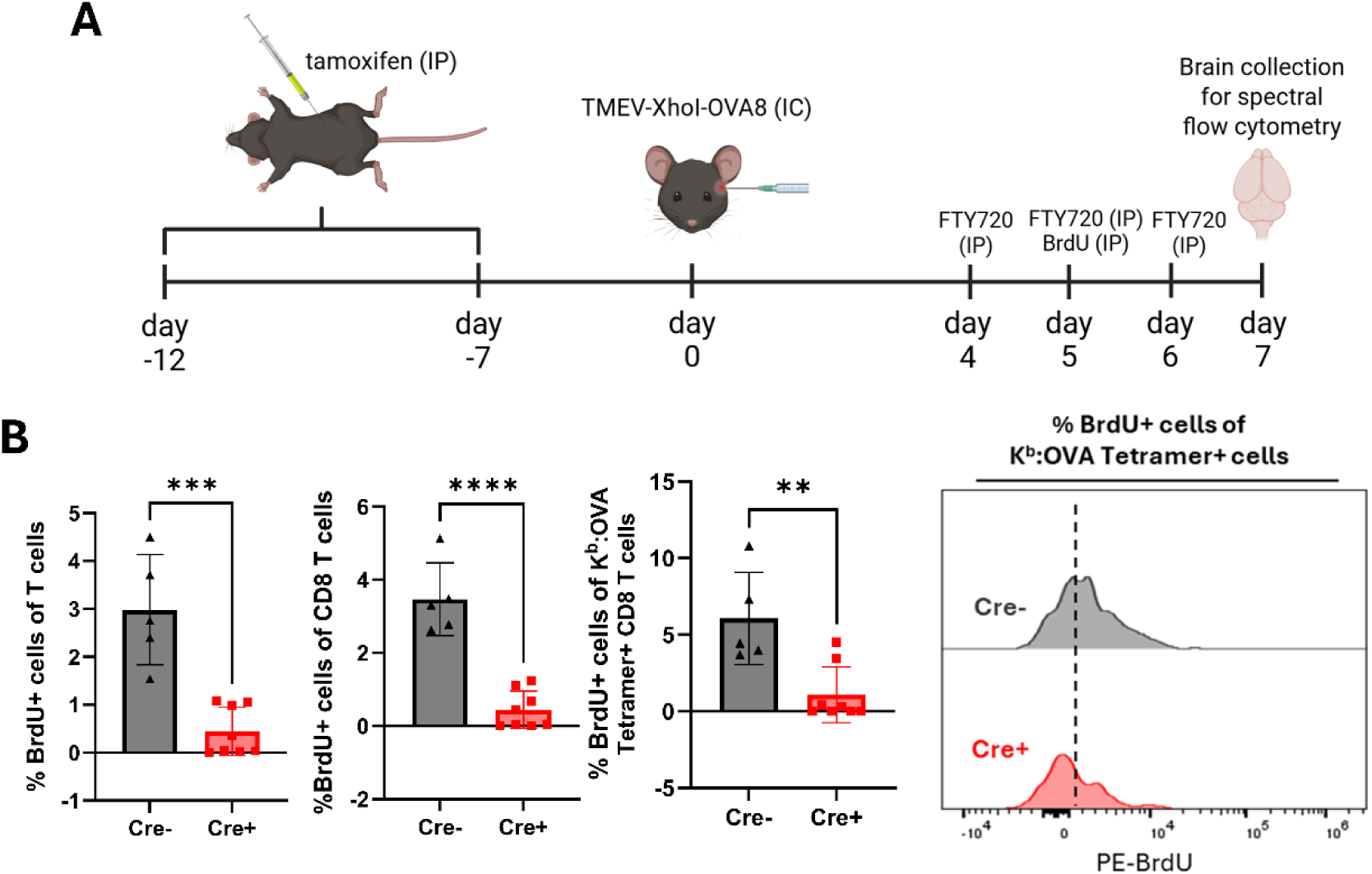
Marked reduction of BrdU labeling on T cells, CD8 T cells and K^b^:OVA Tetramer+ cells is observed in the Tmem119 K^b^ cKO mice compared to cre-littermate controls. (A) Experimental design showing mice were treated with FTY720 or water twice a day starting 4 days after TMEV-OVA infection. 5 days after TMEV-OVA infection, mice received an IP injection of BrdU. Mice were euthanized 7 days after TMEV-OVA infection and perfused. Brains were analyzed using a Cytek Aurora. (B) Bar charts show reduced % BrdU in T cells, CD8 T cells and K^b^:OVA Tetramer+ cells in Tmem119 K^b^ cKO mice compared to cre- littermate controls, as well as representative histograms for Brdu expression in K^b^:OVA Tetramer+ cells. Designation of symbols is as follows: ns or not shown for p > 0.05, * for p ≤ 0.05, ** for p ≤ 0.01, *** for p ≤ 0.001, **** for p ≤ 0.0001. Data presented as mean +/- SD.

### CD8 T cell proliferation induced by microglia H-2K^b^ restricted antigen presentation occurs within the brain

After establishing that H-2K^b^ expression by microglia is required for complete CD8 T cell responses in the brain following neurotropic viral infection, we sought to define the anatomical location of this stimulation. Parabiosis is a powerful method in which two mice are surgically conjoined to share circulation. This approach enables a Tmem119 K^b^ cKO mouse to have a surrogate immune system provided by a Cre- littermate parabiont. We surgically conjoined Tmem119 K^b^ cKO mice to Cre- littermates and waited 3 weeks to ensure shared circulation. At this timepoint, we infected both mice intracranially with TMEV-OVA (Figure 5A). 7 days post TMEV-OVA infection, we PBS perfused both parabionts and analyzed their brain infiltrating immune responses. Following paired wise analysis, we determined that Tmem119 K^b^ cKO mice still exhibited decreased numbers of infiltrating cells and CD8 T cells (Figure 5B) despite sharing peripheral immune system with a Cre- littermate. A trending decline in the numbers of K^b^:OVA Tetramer+ CD8 T cells was also observed. These results demonstrate stimulation of CD8 T cells by H-2K^b^ on microglia occurs within the brain and is not governed by peripheral factors.

**Figure 5.**
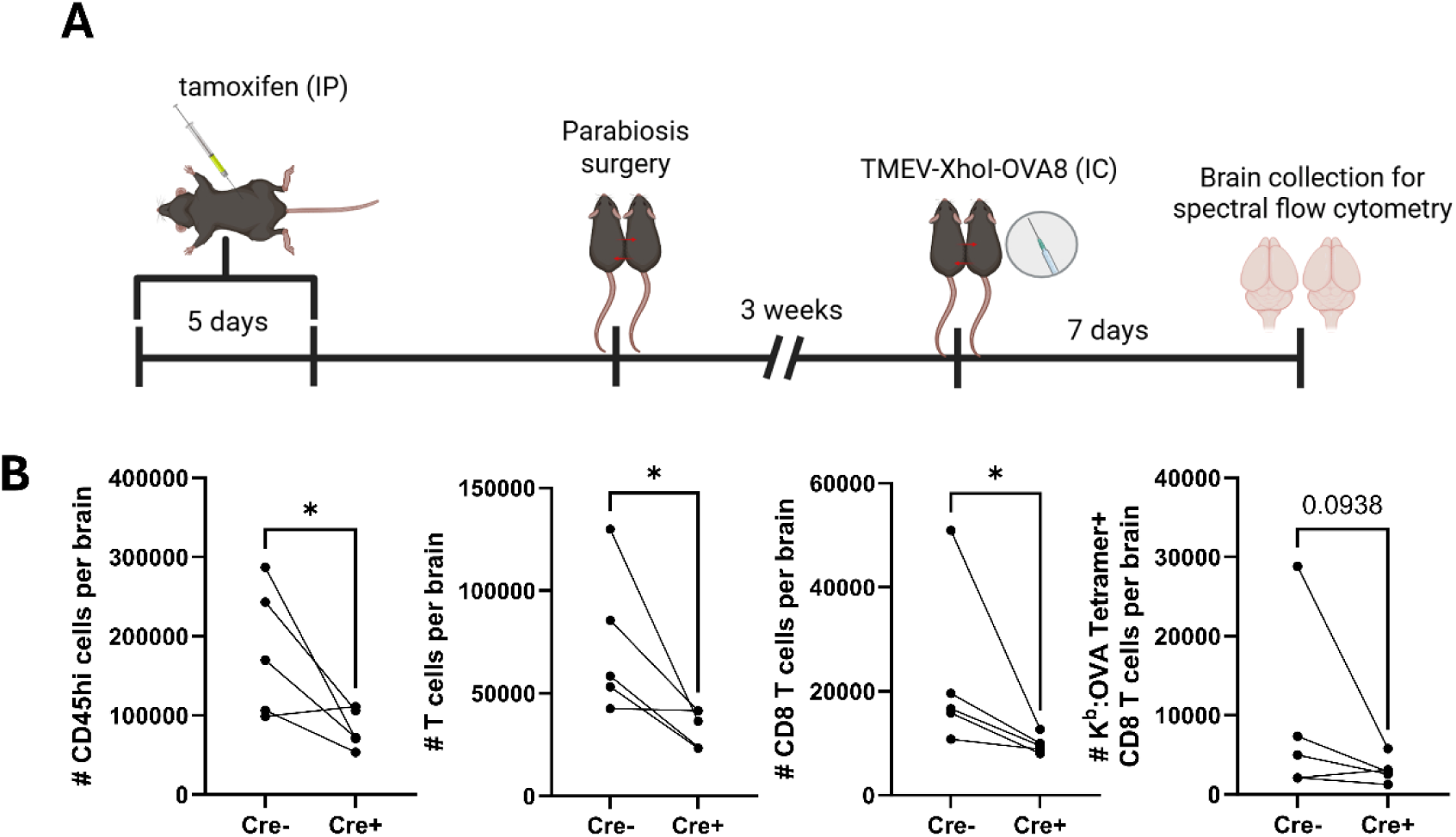
CD8 T cell expansion in the brain is governed by microglia K^b^ restricted antigen presentation and not through peripheral circulating factors. (A) Experimental design showing mice were treated with tamoxifen to activate cre for 5 days, once a day. A Tmem119 K^b^ cKO mouse was then attached to a Cre- littermate through parabiosis surgery. 3 weeks later, mice were infected with TMEV-OVA and euthanized 7 days post infection. Brains were analyzed using spectral flow cytometry. (B) Absolute counts of CD45hi, T cells, CD8 T cells and K^b^:OVA Tetramer+ CD8 T cells per brain. Designation of symbols is as follows: ns or not shown for p > 0.05, * for p ≤ 0.05, ** for p ≤ 0.01, *** for p ≤ 0.001, **** for p ≤ 0.0001. Data presented as mean +/- SD.

### Microglia expression of H-2D^b^ class I molecule enhances CD8 T cell activation and effector status in the brain

During TMEV infection, H-2D^b^ class I molecule presents the immunodominant TMEV peptide, VP2_121-130_, to CD8 T cells, which is required for viral clearance ^17, 18, 30^. We generated a novel Tmem119 D^b^ cKO mice in which H-2D^b^ is deleted from TMEM119+ microglia (Figure 6A). Tmem119 D^b^ cKO mice were infected with TMEV. At 7 days post infection, mice were perfused with 1X PBS and brains were harvested for flow cytometric analysis of CD8 T cell infiltration (Figure 6B). We did not observe changes in frequencies of major immune cell populations, including D^b^:VP2_121-130_ Tetramer+ CD8 T cells (Figure 6C and 6D). However, we observed reduced effector properties in CD8 T cells. This was demonstrated by a marked decrease in perforin protein levels and reduced percentages of perforin+ CD8 T cells and D^b^:VP2_121-130_ Tetramer+ CD8 T cells in 7-day TMEV infected Tmem119 D^b^ cKO mice (Figure 6E). Perforin is a cytolytic protein expressed during the effector stages of antiviral immune responses, orchestrating killing mechanisms essential for complete cytotoxic CD8 T cell function ^24^. This result indicates that H-2D^b^ antigen presentation on microglia are responsible for enhancing CD8 T cell effector status in the brain.

**Figure 6.**
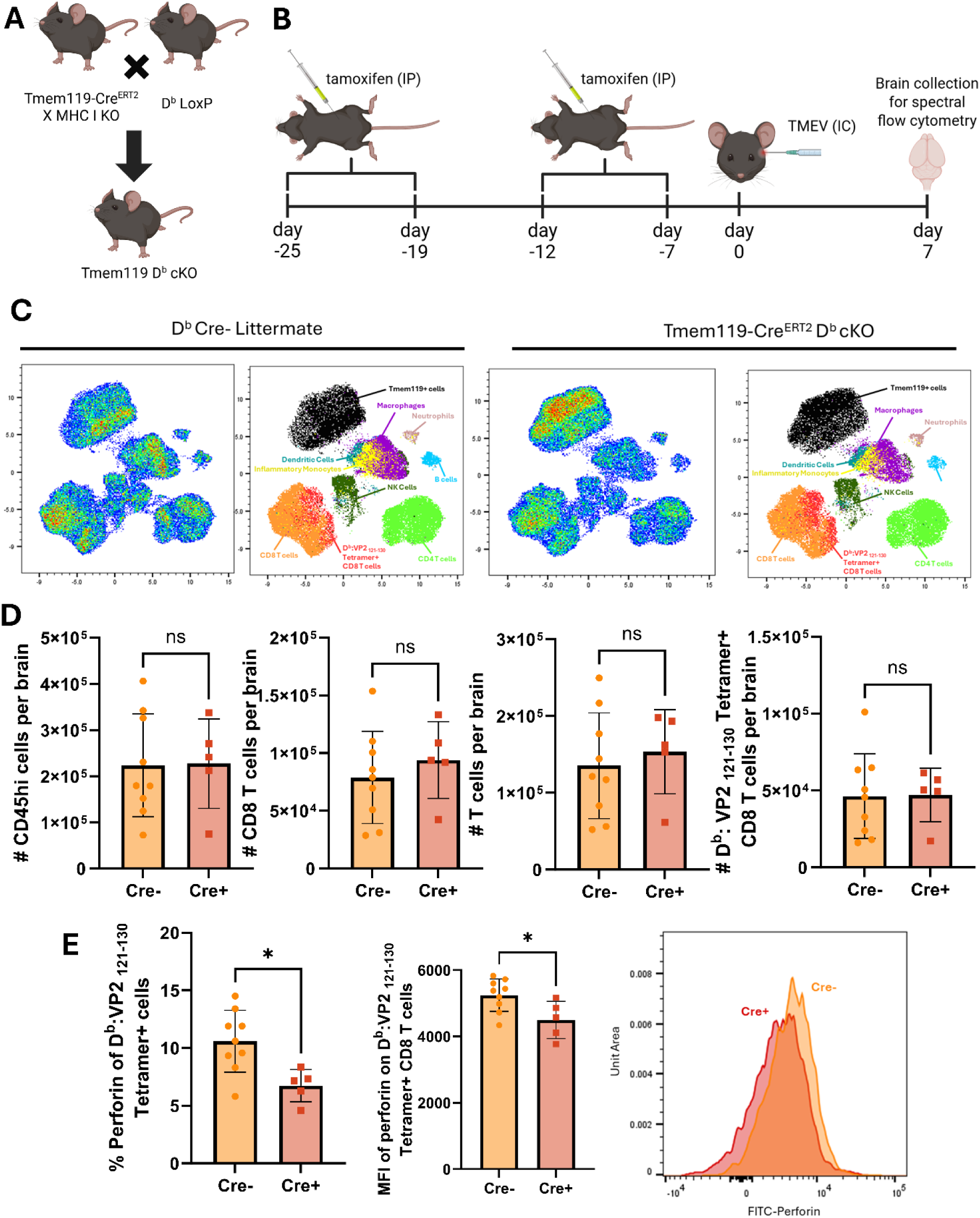
Tmem119 D^b^ cKO mice have decreased perforin expression by CD8 T cells 7 days post TMEV infection. (A) Schematic depicting the Cre-LoxP strategy. Tmem119-Cre^ERT^^2^ mice were crossed to transgenic H-2D^b^ LoxP mice. (B) Experimental design showing mice were treated with tamoxifen to activate cre for 5 days, once a day. After 1 week, another 5 days of tamoxifen injections were employed. Mice were then infected with TMEV and euthanized 7 days post infection. Brains were analyzed using a Cytek Aurora. (C) Representative uniform manifold approximation and projection (UMAP) key, colored to highlight the different cell subsets in the brain for both D^b^ Cre- Littermate and Tmem119-CreER^T^^2^ D^b^ cKO mice. (D) Absolute counts of CD45hi, T cells, CD8 T cells and D^b^:VP2_121-130_ Tetramer+ CD8 T cells per brain are displayed in the bar charts, as well as frequency of perforin on CD8 T cells and D^b^:VP2_121-130_ Tetramer+ cells. (E) Percentages of Perforin+ cells of D^b^:VP2_121-130_ Tetramer+ CD8 T cells are displayed in the bar charts, followed by graph displaying MFI of perforin on of D^b^:VP2_121-130_ Tetramer+ CD8 T cells and a histogram of a representative sample for Perforin expression on D^b^:VP2_121-130_ Tetramer+ CD8 T cells (scaled to Unit Area to normalize by area and not number of events). Designation of symbols is as follows: ns or not shown for p > 0.05, * for p ≤ 0.05, ** for p ≤ 0.01, *** for p ≤ 0.001, **** for p ≤ 0.0001. Data presented as mean +/- SD.

### Microglia expression of D^b^ enhances CD8 T cell mediated blood-brain barrier disruption in a CNS vascular disease model

Our finding that perforin was reduced in CD8 T cells isolated form TMEV infected Tmem119 D^b^ cKO mice was intriguing and prompted us to evaluate these animals for CNS vascular permeability. Our laboratory has determined that perforin expressing CD8 T cells mediate BBB disruption in experimental cerebral malaria and peptide induced fatal syndrome (PIFS) ^24, 25, 31^. PIFS is a model generated in our laboratory that consists of intravenously injecting the viral immunodominant peptide VP2_121-130_ in mice at day 7 post TMEV infection (Figure 7A) ^32^. VP2_121-130_ peptide administration stimulates activated D^b^:VP2_121-130_ epitope specific CD8 T cells which in turn induce severe blood-brain barrier (BBB) disruption within 4 hours and is ultimately fatal at 24 hours ^33^. This BBB disruption is defined by the loss of cerebrovascular endothelial cell tight junction integrity. and is governed by perforin expressing CD8 T cell response, consistent with our findings in experimental cerebral malaria ^24, 31^ ^33^. Therefore, we investigated the impact of differential expression of perforin by D^b^:VP2_121-130_ Tetramer+ CD8 T cells on BBB disruption in Tmem119 D^b^ cKO mice using the PIFS model. We noticed a marked reduction in overall numbers of infiltrating blood-derived cells (CD45hi cells), total T cells, CD8 T cells and D^b^:VP2_121-130_ Tetramer+ CD8 T cells, indicating that the loss of H-2D^b^ on microglia attenuates the immune response in the brain (Figure 7B). To assess BBB integrity, in a separate experiment, we intravenously injected FITC-albumin in both Tmem119 D^b^ cKO mice and Cre- littermate controls 1 hour before euthanasia (Figure 7C). We observed reduced CNS vascular permeability as demonstrated by FITC-albumin leakage into brain parenchyma in sagittal cut sections of Tmem119 D^b^ cKO mouse brains when compared to Cre- littermates (Figure 7D). Compromised BBB integrity in Cre-controls was therefore consistent with heightened levels of perforin expression by D^b^:VP2_121-130_ Tetramer+ CD8 T cells (Figure 6E and 7D). In contrast, Tmem119 D^b^ cKO mice have preserved vascular integrity due to the absence of H-2D^b^ restricted antigen presentation by microglia in the PIFS model which is also consistent with CD8 T cells having markedly reduced perforin expression (Figure 6E and 7D).

**Figure 7.**
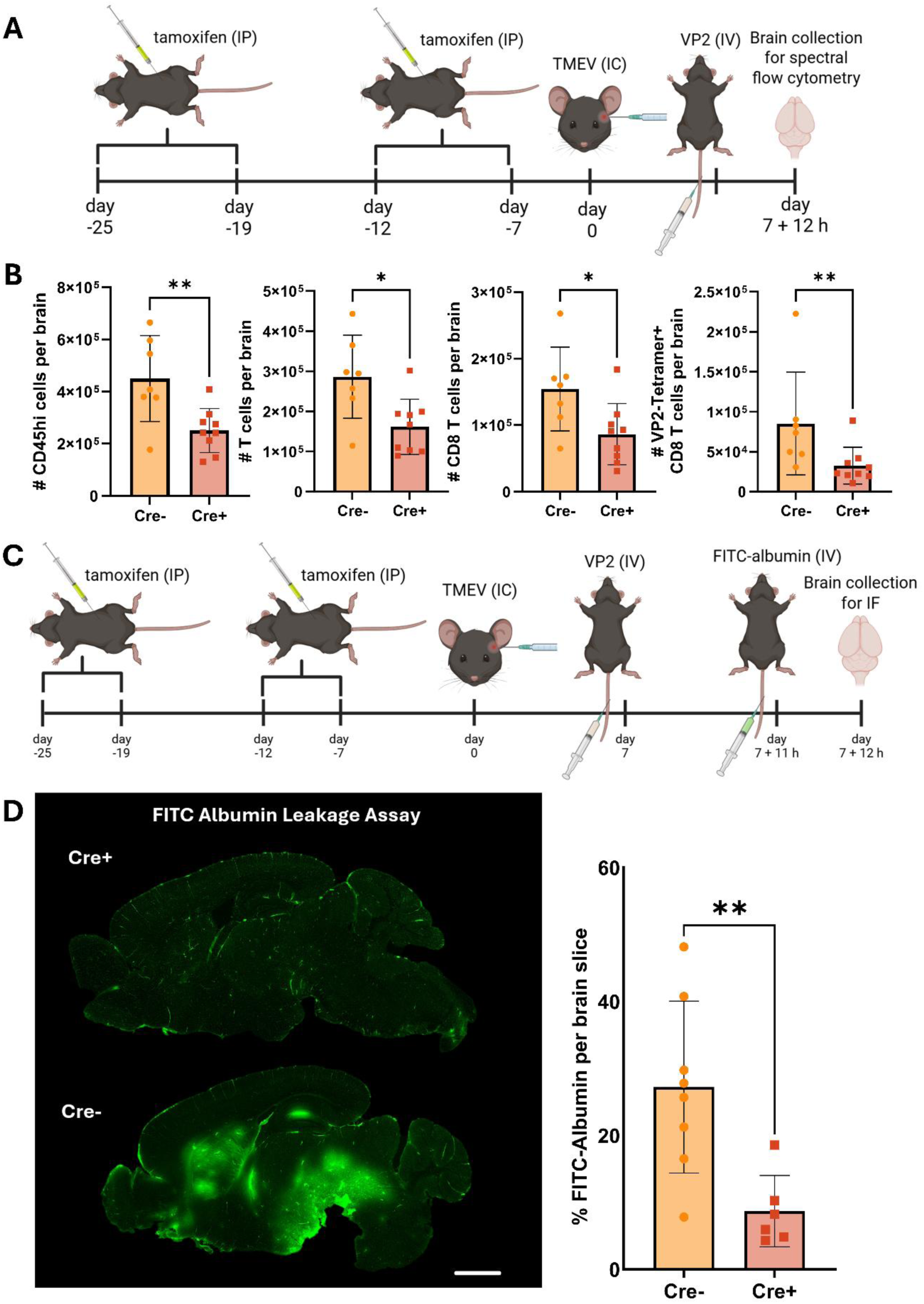
Peptide-induced fatal syndrome elicits increased cell infiltration in comparison to Tmem119 D^b^ cKO mice. (A) Mice were intracranially infected with TMEV. 7 days later, mice received the TMEV immunodominant peptide VP2_121–130_ intravenously. Injection of VP2121–130 peptide will induce lethal blood-brain barrier. 12 hours post injection of VP2_121–130_, mice were euthanized, perfused, and analyzed. (B) Absolute counts of CD45hi, T cells, CD8 T cells and D^b^:VP2_121-130_ Tetramer+ CD8 T cells per brain are displayed in the bar charts. (C) Mice were intracranially infected with TMEV. 7 days later, mice received the TMEV immunodominant peptide VP2_121–130_ intravenously. 11 hours after the VP2_121–130_ injection, FITC-albumin was administered intravenously. 12 hours post injection of VP2_121–130_, mice were euthanized and analyzed. (D) Representative sagittal images of FITC-albumin injected mice 12 hours post VP2_121–130_ injection, displaying BBB leakage differences between Cre+ and Cre- groups. Quantification is displayed on the left. Scale bar = 2 mm. Designation of symbols is as follows: ns or not shown for p > 0.05, * for p ≤ 0.05, ** for p ≤ 0.01, *** for p ≤ 0.001, **** for p ≤ 0.0001. Data presented as mean +/- SD.

### CD8 T cell-mediated BBB disruption is exacerbated by microglia D^b^ restricted antigen presentation and is not impacted by peripheral circulating factors

To determine the location of CD8 T cell stimulation by MHC class I expressing microglia, we performed parabiosis surgery to conjoin Tmem119 D^b^ cKO mice to Cre- littermate controls. 3 weeks post-surgery, we intracranially infected both mice with TMEV. At 7 days post infection, we i.v. administered VP2_121-130_ peptide intravenously in both mice to induce PIFS. 11 hours post-VP2_121-130_ peptide delivery and 1 hour prior to euthanasia, all mice were i.v. injected with FITC-albumin to measure BBB disruption (Figure 8A). Despite sharing circulation to a Cre- littermate control, Tmem119 D^b^ cKO mice presented with greatly reduced BBB disruption (Figure 8B). Paired wise quantification of sagittal cuts obtained from 4 sets of parabionts were analyzed (Figure 8C). This data demonstrates that shared circulation and peripheral factors do not restore CD8 T cell mediated BBB disruption when microglia was no longer able to present antigen via H-2D^b^.

**Figure 8.**
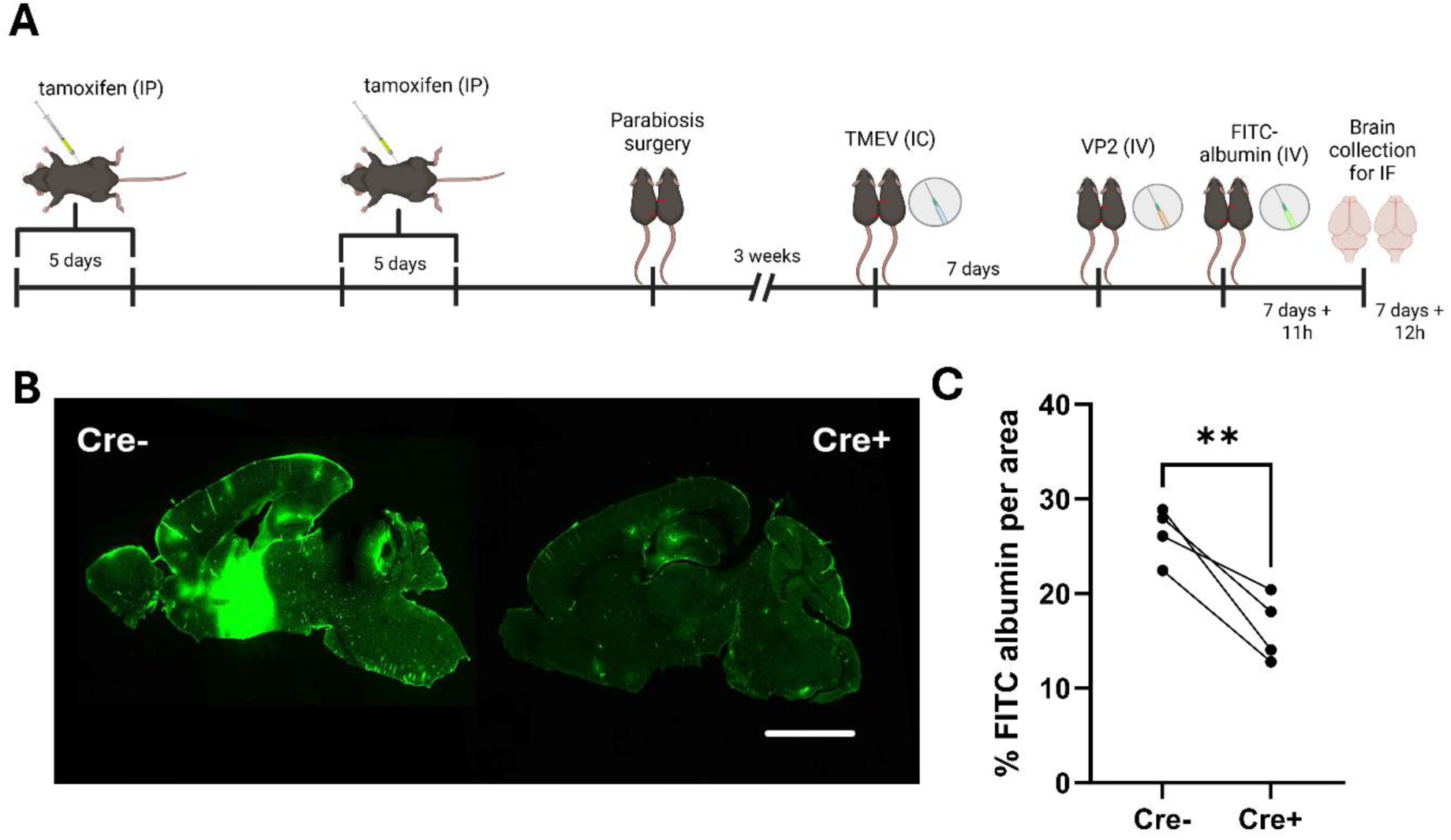
Tmem119 D^b^ cKO mice are protected from BBB disruption despite sharing circulation with a Cre- littermate parabiont. (A) Experimental design showing mice were treated with tamoxifen to activate cre for 5 days, once a day, and then another 5 days 1 week later. A Tmem119 D^b^ cKO mouse was then attached to a Cre- littermate through parabiosis surgery. 3 weeks later, mice were infected with TMEV. 7 days later, mice received the TMEV immunodominant peptide VP2_121–130_ intravenously. 11 hours after the VP2_121–130_ injection, FITC-albumin was administered intravenously. 12 hours post injection of VP2_121–130_, mice were euthanized and analyzed. (B) Representative sagittal images of FITC-albumin injected mice 12 hours post VP2_121–130_ injection, displaying BBB leakage differences between Cre+ and Cre- groups. Quantification is displayed on the left. Scale bar = 2 mm. Designation of symbols is as follows: ns or not shown for p > 0.05, * for p ≤ 0.05, ** for p ≤ 0.01, *** for p ≤ 0.001, **** for p ≤ 0.0001. Data presented as mean +/- SD.

### Microglia MHC class I antigen presentation does not affect the generation of memory CD8 T cells

TMEV infection is cleared from the brain approximately between 14 - 21 days post infection ^8, 19^. It has been recently shown that TMEV infection results in the accumulation and retention of antigen specific CD8 TRM cells within the brain at 30 days post infection ^3^. However, it is not known if microglial MHC class I responses are involved in the generation of CD8 TRM cells in the brain. To define the contribution of H-2D^b^ on microglia for CD8 T cell memory formation in the brain, we infected both Tmem119 D^b^ cKO mice and Cre- littermate controls and analyzed the numbers of CD8 TRM cells 30 days post infection (Figure 9A). We did not observe differences between the groups at this time point, as the number of D^b^:VP2_121-130_ Tetramer+ CD8 T cells was similar in Tmem119 D^b^ cKO and Cre- mice (Figure 9B). CD69+ CD103+ of D^b^:VP2_121-130_ Tetramer+ cells were also not altered (data not shown).

**Figure 9.**
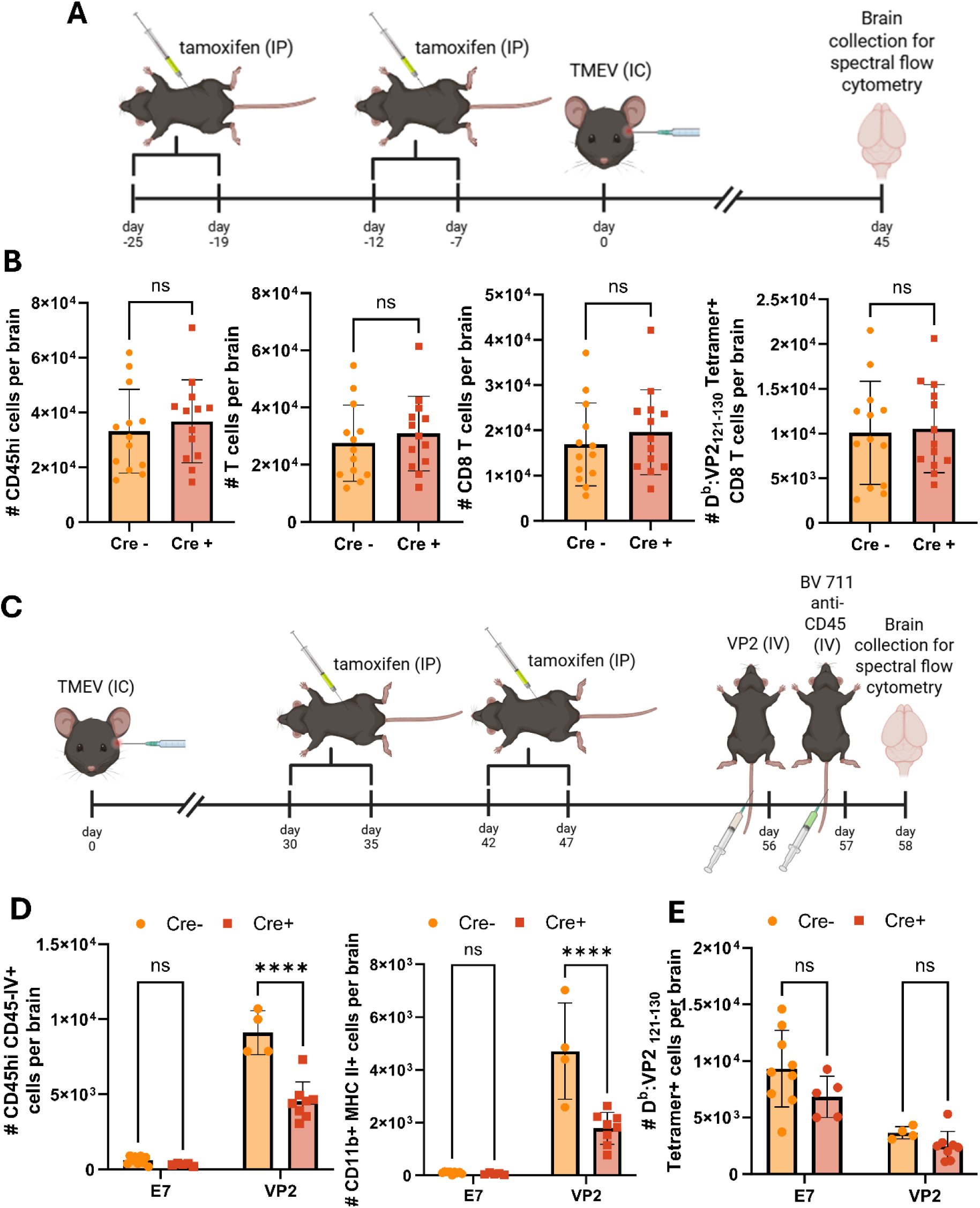
Microglia D^b^ restricted antigen presentation is not required for CD8 TRM cell formation in comparison to Cre- littermates. However, upon secondary challenge, Tmem119 D^b^ cKO mice have attenuated immune cell infiltration. (A) Experimental design showing mice were treated with tamoxifen to activate cre recombinase for 5 days, once a day, and then another 5 days 1 week later. Mice were intracranially infected with TMEV and allowed 45 days for resolution of the infection and TRM generation. Mice were then euthanized, perfused, and analyzed. (B) Absolute counts of CD45hi, T cells, CD8 T cells and D^b^:VP2_121-130_ Tetramer+ CD8 T cells per brain are displayed in the bar charts. (C) Mice were intracranially infected with TMEV and allowed 30 days for resolution of the infection and TRM generation. Later, mice were treated with tamoxifen to activate cre recombinase for 5 days, once a day, and then another 5 days 1 week later. Following tamoxifen injections, mice received either the TMEV immunodominant peptide VP2_121–130_, or irrelevant peptide E7 two days before euthanasia. Injection of VP2_121–130_ peptide will reactivate TMEV specific T cells including TRMs in the brain. 24 hours post VP2_121–130_ injections mice received an intravenous injection of BV711 anti-CD45 conjugated antibody to label CD45+ cells within the blood circulation. 24 hours post injection of antibodies, mice were euthanized, perfused, and analyzed. (D) Absolute counts of infiltrating cells (CD45hi CD45-IV+ cells per brain) and myeloid cells (CD11b+ MHC II+ cells per brain). (E) Absolute counts of D^b^: VP2_121-130_ Tetramer+ CD8 T cells per brain. Designation of symbols is as follows: ns or not shown for p > 0.05, * for p ≤ 0.05, ** for p ≤ 0.01, *** for p ≤ 0.001, **** for p ≤ 0.0001. Data presented as mean +/- SD.

Our laboratory has also developed a model to study the reactivation of CD8 TRM cells through administration of immunodominant viral peptide, VP2_121-130_, post viral clearance ^3^. This stimulation results in the reactivation of D^b^:VP2_121-130_ epitope specific CD8 TRM cells and sublethal BBB disruption. To assess the contribution of H-2D^b^ antigen presentation on microglia for reactivation of antigen specific CD8 TRM cells, immune cell infiltration in general, and BBB disruption, we infected mice with TMEV and then injected tamoxifen to activate cre recombinase (Figure 9C). This allows us to investigate the effects of H-2D^b^ expression on microglia at the time of the reactivation. Anti-CD45 antibody i.v. labeling was used to identify cells residing in vasculature verses brain parenchyma. Animals were perfused to ensure that immune cells in vasculature were excluded from analysis. 48 hours after VP2_121-130_ or control E7 peptide, we observed an increase in the numbers of infiltrating cells in both Tmem119 D^b^ cKO mice and Cre-littermate controls (Figure 9D). However, Tmem119 D^b^ cKO mice that received VP2_121-130_ injection exhibited a decrease in the immune cell infiltration in comparison to Cre-littermates that also received VP2_121-130_ (Figure 9D). Interestingly, in agreement with previous data from our laboratory, we observed an expansion of CD11b+ MHC II+ cells following VP2_121-130_ injection ^3^. This population was attenuated in the Tmem119 D^b^ cKO mice, suggesting that microglia MHC class I is involved in the recruitment of overall infiltrating cells and the expansion of a myeloid cell population, potentially following engagement by CD8 T cell. Although we did find altered immune cell infiltration overall, we did not observe differences in the numbers of D^b^:VP2_121-130_ Tetramer+ CD8 T cells between Tmem119 D^b^ cKO and Cre- littermate controls in the numbers of (Figure 9E).

### Microglial MHC class I contributes to the expansion of CD8 TRM cell populations in the brain during antigen recall response

To assess the contribution of microglia MHC class I in the recall response of CD8 TRM cells, we TMEV infected Tmem119 D^b^ cKO mice and Cre- littermate controls before tamoxifen injections. D^b^ expressing mice resolve TMEV infection in 14-21 days ^8, 19^. We then waited until day 30 to administer tamoxifen to inactivate D^b^ class I molecule in microglia (Figure 10A). FTY720 was administered 3 days prior VP2_121-120_ i.v injection and 2 days after to restimulate memory D^b^:VP2_121-130_ epitope specific CD8 TRM cells (Figure 10A). This approach allows blockade of peripheral lymphocyte trafficking into the brain including peripheral memory CD8 T cells, thereby isolating D^b^:VP2_121-130_ epitope specific CD8 TRM cell responses. First, we confirmed that FTY720 treatment resulted in decreased T cell frequencies in the blood (Figure S3). Meanwhile, through analysis of the brain we identified that Tmem119 D^b^ cKO mice had a marked decrease in both frequencies and numbers of D^b^:VP2_121-130_ Tetramer+ CD8 T cells, accompanied by a decrease in both frequencies and numbers of total infiltrating cells (CD45hi+ cells) and myeloid cells (CD11b+ MHC II+ cells) (Figure 10B, C and D). Importantly, we observed reduced BBB permeability through reduced gadolinium leakage into the tissue in Tmem119 D^b^ cKO mice (Figure 10E). It is important to note that both Tmem119 D^b^ cKO mice and Cre- littermate controls received FTY720 and VP2_121-120_ injections. Therefore, reduced D^b^:VP2_121-130_ Tetramer+ cell expansion in the brain and reduced BBB permeability is the result of microglia H-2D^b^ antigen presentation being conditionally ablated. This demonstrates a critical requirement of microglia D^b^ restricted antigen presentation in the recall CD8 TRM response to the immunodominant D^b^:VP2_121-130_ epitope during neurotropic virus infection.

**Figure 10.**
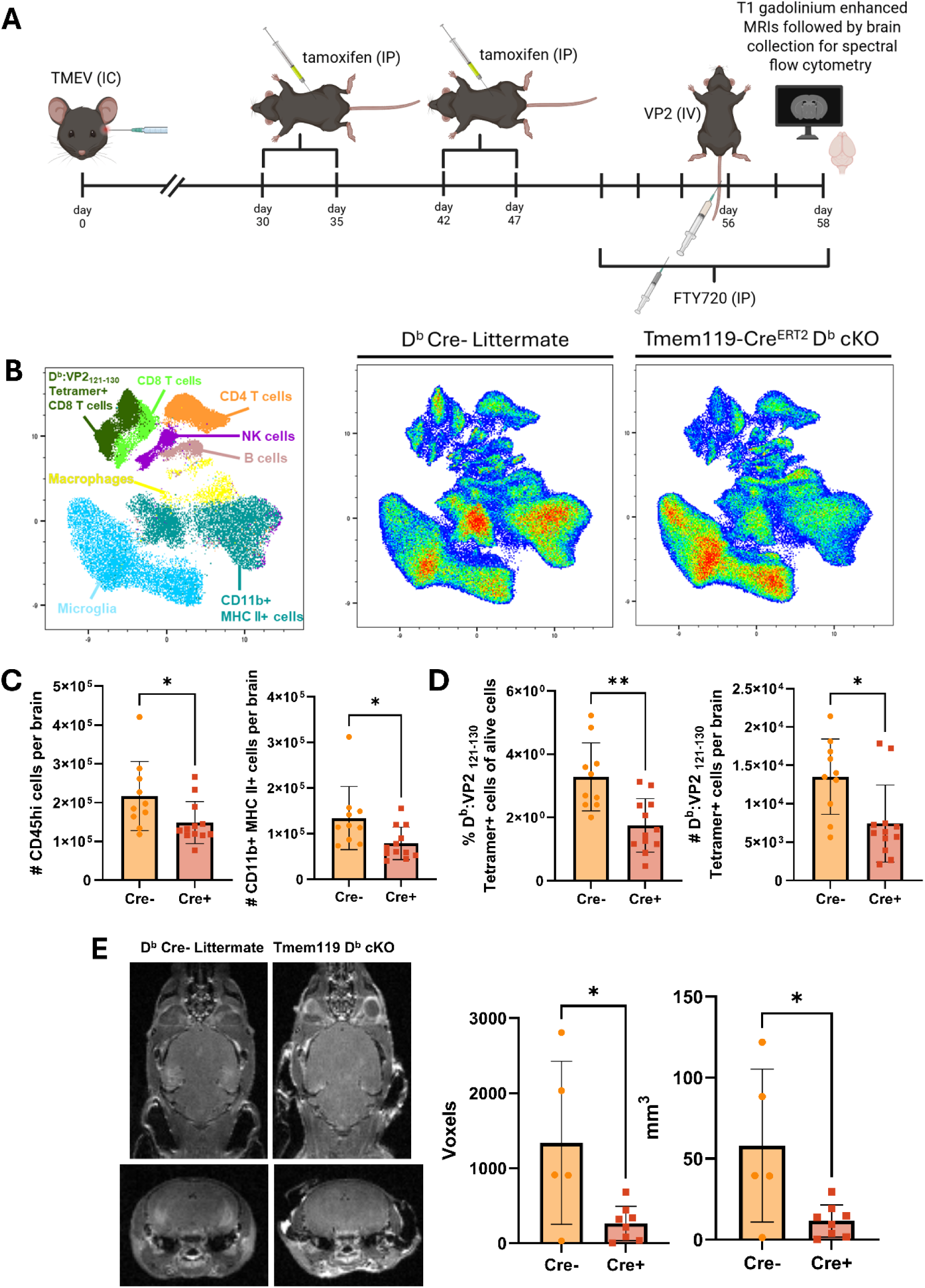
CD8 TRM cell responses are decreased in Tmem119 D^b^ cKO mice and accompanied by reduced numbers of infiltrating cells and BBB permeability. (A) Experimental design showing mice were intracranially infected with TMEV and allowed 30 days for resolution of the infection and TRM generation. Later, mice were treated with tamoxifen to activate cre recombinase for 5 days, once a day, and then another 5 days 1 week later. Following tamoxifen injections, mice received an intravenous injection of the TMEV immunodominant peptide VP2_121–130_ to reactivate TMEV specific T cells including CD8 TRMs in the brain. 3 days before and 1 day after VP2_121–130_ injections, mice received intraperitoneal injections of FTY720 twice a day. 48 hours post VP2_121–130_ injections, mice were imaged using T1 gadolinium enganced MRIs and then euthanized, perfused, and analyzed. (B) UMAPs displaying frequencies of alive of several immune cell populations are displayed. (C) Absolute counts of infiltrating cells (#CD45hi+ cells) and myeloid cells (#CD11b+ MHC II+ cells) per brain, as well as counts and numbers of D^b^:VP2_121-130_ Tetramer+ cells are displayed. (D) BBB permeability was measured by gadolinium enhanced MRIs analysis using T1 weighted MRIs. Representative images are displayed on the left followed by quantification on the right. Designation of symbols is as follows: ns or not shown for p > 0.05, * for p ≤ 0.05, ** for p ≤ 0.01, *** for p ≤ 0.001, **** for p ≤ 0.0001. Data presented as mean +/- SD.

## Discussion

In this study, we capitalized on the magnitude of immunodominant CD8 T cell responses generated against neurotropic TMEV infection to define the impact of discrete MHC class I molecule restricted antigen presentation by microglia. Through our ability to conditionally inactivate H-2K^b^ and H-2D^b^, our study was able to define the contribution of individual microglia MHC class I antigen presentation during acute and memory responses. We defined that H-2K^b^ antigen presentation on microglia is essential for the expansion of K^b^:OVA Tetramer+ CD8 T cells during neurotropic virus infection. Our data specifies that this expansion is due to proliferation of CD8 T cells and occurs as a consequence of microglia H-2K^b^ restricted antigen presentation. Furthermore, we define that H-2D^b^ antigen presentation on microglia is important for heightened activation of antiviral D^b^:VP2_121-130_ epitope specific CD8 T cells as observed by enhanced perforin expression. This modulation results in lower overall immune cell infiltration and reduced BBB disruption in the PIFS model. Importantly, these effects are localized to the CNS. Finally, post clearance of the TMEV virus infection, we demonstrate microglia H-2D^b^ is required for complete immune cell infiltration during a secondary encounter with virus antigen. CD8 TRM cells are activated by H-2D^b^ restricted antigen presentation by microglia, resulting in increased immune cell infiltration.

CD8 T cell expansion in the brain is a key element to many neurological diseases ^34^. To our knowledge, this work is the first demonstration of MHC class I antigen presentation by microglia directly activating CD8 T cells to promote BBB disruption. Furthermore, we dissected the contribution of discrete MHC class I molecules on microglia in shaping CD8 T cell response in the brain.

Historically, whole cell ablation strategies have been employed to define the contribution of microglia to neuroinflammation, including the influence on adaptive immune cell responses. Pexidartinib (PLX3397) is a CSF1 receptor inhibitor widely used for microglia depletion. However, this depletion strategy is not limited to microglia or even CNS myeloid cells in general ^20^. Previous studies have shown that PLX3397 administration decreases splenocyte numbers and depletes CD11c+ cells, implying peripheral T cell priming by dendritic cells would be impaired ^8^. Additional investigation of PLX3397 administration revealed circulating and bone marrow derived immune cells beyond the mononuclear phagocyte system were also affected, suggesting the need for caution when interpreting results employing this depletion strategy ^20^. Recently, an alternative approach using PLX5622 has been proposed to be more microglia specific. However, PLX5622 also has significant non-microglia off target effects that could contribute to results observed employing this drug ^21, 35^. Importantly, whole cell depletion of microglia nullifies the ability to study discrete microglial functions in experimental neurologic disease models.

Aside from cellular depletion strategies, CX3CR1 Cre expressing mice are commonly used to conduct gene ablation in microglia. Previous work employing this Cre strategy has been important in highlighting microglia as an antigen presenting cell, capable of modulating CD8 T cell responses in the brain ^8^. During neurotropic viral infection, H-2D^b^ antigen presentation by microglia was put forward as a major regulator for CD8 T cell response and brain atrophy as a long-term consequence ^8^. However, studies employing CX3CR1 promoter induced green fluorescent protein (GFP) revealed monocytes, NK cells, circulating and skin resident dendritic cells were also impacted in addition to microglia ^22, 26^. In tamoxifen-induced systems, waiting a 6-week interval after Cre recombinase activation allows peripheral CX3CR1 expressing cells in the periphery to turnover, reducing the off-target effects of this approach ^8^. However, brain resident CX3CR1 expressing cells such as long-lived CNS residing perivascular macrophages remain impacted. Therefore, while the CX3CR1 Cre line has been commonly used as a microglia-specific strategy, the expression of CX3CR1 by non-microglial cells in the brain still needs to be addressed when interpreting results obtained with this strategy. The Tmem119-Cre^ERT^^2^ system employed in this study is a more specific approach to assess microglial function ^27, 28^. Tmem119 is a transmembrane protein that is highly expressed by microglia in both mice and humans. Importantly, the Tmem119-EGFP system was found to target parenchymal microglia, but not other brain macrophages and blood monocytes ^27^.

Microglia MHC class II antigen presentation has been investigated through analysis of human tissue and experiments using preclinical models. MHC class II expressing microglia have been reported in brains of patients with Parkinson’s disease (PD) and Alzheimer’s disease (AD) ^14^. It has been reported in a mouse model that the expression of full-length human α-synuclein increases the expression of MHC II by microglia ^36^. Deletion of MHC II significantly reduced microgliosis following AAV2-SYN induction in this PD model ^36^. However, in studies using experimental autoimmune encephalomyelitis (EAE) and cuprizone-induced demyelination models, deletion of MHC II in CX3CR1+ cells did not alter disease onset, progression, or severity ^37^. Much less studied is the extent MHC class I antigen presentation by microglia shapes immune responses and neuropathology, especially regarding the use of precision tools that specifically target microglia ^38^. Defining the role of MHC class I responses in microglia is of critical need, given microglia have been defined as a major cellular source of MHC class I expression in the CNS of both mice and humans when compared to other brain resident cell types ^13^. In this study, we therefore employed Tmem119-Cre^ERT^^2^ mice crossed with H-2K^b^ or H-2D^b fl/fl^ mice. This approach allowed us to investigate the role of discrete MHC class I molecules in TMEM119+ microglia, other than assessing MHC class I function as a singular molecule.

Strategies to conditionally ablate β2-microglobulin expression have been employed to understand MHC class I biology MHC class I function *in vivo*. However, this is a very broad knockout strategy. β2-microglobulin stabilizes hundreds of protein complexes as diverse as Hfe, FcRn, MICA and MICB, Qa-1, MR1, TL antigens, and CD1 ^39, 40, 41, 42, 43, 44, 45^. These numerous protein species govern major branches of inflammation including iron metabolism, antibody transport to the brain, innate immune recognition of stressed and apoptotic cells, adaptive immune cell function, NK cell killing, and NK T cell activation. Off target effects of removing these major drivers of neuroinflammation confounds interpretation of results obtained using targeted ablation of β2m ^40, 46^. Furthermore, the highly polymorphic mouse H-2 and human HLA genes are far from redundant in function ^47^. In the TMEV infection model alone, H-2 genes govern virus persistence and susceptibility to demyelination, with D^b^ being genetically defined as a critical protecting allele ^17^. Furthermore, in experimental cerebral malaria (ECM), K^b^-restricted antigen engagement on brain endothelium was shown to be pivotal for CD8 T cell vascular adhesion, arrest, infiltration and activation. Meanwhile, only CD8 T cell receptor recognition of D^b^-restricted antigens resulted in killing of mouse brain microvascular endothelial cell and subsequent neuronal cell death, supporting that K^b^ and D^b^ regulate distinct patterns of disease onset ^48^. Finally, notable HLA genes have been associated historically with resistance to human pathogens ^49, 50, 51^. Our data supports the heterogeneity behind MHC class I molecules as Tmem119 K^b^ cKO mice exhibit a decreased antigen specific CD8 T cell response following a neurotropic virus infection and Tmem119 D^b^ cKO mice do not follow the same pattern. In contrast, Tmem119 D^b^ cKO mice exhibit decreased CD8 T cell activation observed by reduced perforin expression, while Tmem119 K^b^ cKO mice did not result in this same phenotype, supporting the importance of conditional ablation of discrete MHC I genes in designated cell types.

The reduction of K^b^:OVA Tetramer+ CD8 T cells numbers observed in TMEV-OVA infected Tmem119 K^b^ cKO mice used in this study is likely due to a stimulatory event taking place in the brain parenchyma. The Daniel strain of TMEV is a neurotropic picornavirus that does not directly infect microglia yet results in a robust brain infiltrating CD8 T cell response ^16^. Moreover, incorporating OVA antigen sequence into recombinant TMEV provides an important tool to also study H-2K^b^ restricted CD8 T cell responses in the brain ^23^. It is important to note that although the insertion of OVA to the TMEV virus facilitates the tracking of antigen specific response, it does not alter the course of infection Our study further explains the importance of microglia class I molecules on CD8 T cell proliferation and effector function in the brain. While it is known that CD11c+ APCs are critical for early priming of CD8 T cells against the immunodominant TMEV peptide VP2_121-130_, it was not fully understood whether other NVU cell types could provide a secondary stimulatory event that expands the antigen specific CD8 T cell response in the brain parenchyma ^19^. Our data supports a working model that after TMEV infection, antigen-specific CD8 T cells are generated peripherally before infiltrating the brain to engage H-2K^b^ on microglia. The experimental design employing injections of FTY720 at day 4 post infection allowed the first few CD8 T cells to infiltrate and blocked further infiltration from day 4 until day 7. That, combined with BrdU labeling showed that K^b^ restricted antigen presentation on microglia leads to increased cell proliferation of early infiltrating of K^b^:OVA Tetramer+ CD8 T cells, partially explaining the decreased numbers of K^b^:OVA Tetramer+ CD8 T cells found at 7 days post infection in Tmem119 K^b^ cKO mice. Although we observed that cell proliferation is decreased in CD8 T cells of Tmem119 K^b^ cKO mice, it is possible that this stimulatory signal can also lead to regulation of overall immune cell infiltration. This is supported by our observed decrease in other immune cell populations including inflammatory monocytes and macrophages. Previous work from our laboratory has suggested that brain resident CD8 T cells can trigger an immune response that leads to enhanced myeloid cell infiltration and microglial activation ^3^. Finally, our employment of parabiosis techniques in this study confirms that differential peripheral priming of CD8 T cells is not affecting the observed neuroinflammatory phenotype in mice with MHC class I deletion in microglia.

Perforin expression was found to be decreased in Tmem119 D^b^ cKO mice. Although the numbers of D^b^:VP2_121-130_ Tetramer+ CD8 T cells were not different in the Tmem119 D^b^ cKO mice, we observed a marked reduction in the percentages of perforin+ D^b^:VP2_121-130_ Tetramer+ CD8 T cells as well as a reduction in protein expression visualized by MFI values. Perforin is a key cytolytic protein produced by CD8 T cells. This pore-forming protein is critical for cell death and viral clearance during infectious diseases, and it has been defined as a key mechanism for BBB disruption in ECM and the PIFS model ^24, 25^. In the Tmem119 D^b^ cKO, the interaction between infiltrating D^b^:VP2_121-130_ Tetramer+ CD8 T cells might be a key step in further activating these CD8 T cells and not increasing its proliferation as seen in the Tmem119 K^b^ cKO mice. We therefore determined if modulation of CD8 T cell perforin by microglia H-2D^b^ would affect neurotropic viral infection. The PIFS model provides means to assess modulatory effects of perforin on BBB disruption including gene dosage ^24^. Perforin-heterozygous mice have reduced BBB disruption as compared to wild type C57BL/6 mice in the PIFS model ^24^. Similarly, we have found that Tmem119 D^b^ cKO mice with their reduced perforin expression by D^b^:VP2 epitope specific CD8 T cells have attenuated immune cell infiltration and BBB disruption. We contend that the stimulatory effect of CD8 T cells provided by H-2D^b^ is further activating D^b^:VP2_121-130_ epitope specific CD8 T cells, increasing their perforin production and capacity to induce BBB disruption.

Microglia have been linked to BBB disruption in numerous contexts ^52, 53^. Although microglia are not anatomically described as being tightly opposed with endothelial cells at the NVU, there are reports of ramified populations residing in close proximity to brain capillaries ^54^. Furthermore, microglia respond to BBB disruption events through expression of inflammatory mediators plus debris removal to promote BBB restoration ^55^. The upregulation of MHC I is part of this inflammatory response although it happens regardless of BBB disruption ^8, 13^. Interestingly, *in vivo* imaging experiments defined that brain resident microglia migrate to the cerebral vasculature during systemic inflammation ^56^. We hypothesize that the upregulation of MHC class I molecules together with the migration of microglia to areas that surround the cerebral vasculature might position microglia at an optimal location for controlling CD8 T cell activation and further exacerbating BBB disruption. Importantly, this modulation was also confirmed to be brain-centered as observed by parabiosis experiments.

The mechanisms underlying microglial antigen presentation are still unknown. Historically, the concept of cross-presentation has been restricted to dendritic cells which engulf cellular debris and present processed peptides on MHC class I molecules to CD8 T cells ^57^. However, it has also been reported that brain endothelial cells can cross-present during Plasmodium berghei ANKA infection ^58^. In this study, *in vitro* and *in vivo* imaging methodology was employed to observe endothelial cell cross presentation of OVA antigen ^58^. This resulted in clear tracking of antigen processing and presentation in culture and also the arrest of T cells engaging brain vasculature during acute infection ^58^. Therefore, it is possible that additional NVU cell types can cross-present antigens. More specifically, microglia could be engulfing cellular debris from TMEV infected neurons and presenting viral antigen on MHC class I molecules to stimulate brain infiltrating CD8 T cells. Additionally, in our study, the modulation of perforin expression on D^b^:VP2_121-130_ Tetramer+ CD8 T cells by MHC class I on microglia potentiates the BBB damage. It is unclear whether this regulation would be beneficial in different disease models with different degrees of BBB disruption.

Adaptive immune cell memory is an essential protective mechanism against recurring infectious diseases. TMEV infection generates memory D^b^:VP2_121-130_ epitope specific CD8 T cells which reside in the brain long after viral clearance ^3^. Importantly, our data demonstrates that the microglia H-2D^b^ expression is not required for maintenance of these memory CD8 T cells in the brain ^3^. However, upon secondary encounter with the immunodominant viral peptide VP2_121-130_, ablation of H-2D^b^ in microglia results in reduced immune cell infiltration from the blood to the brain parenchyma. Although at first we did not see a decrease in the numbers of D^b^:VP2_121-130_ Tetramer+ CD8 T cells in the Tmem119 D^b^ cKO mice, when isolating the CD8 TRM response using FTY720 treatment, we observed an important interaction between H-2D^b^ expressing microglia and CD8 TRM cells in the brain. This decreased expansion of CD8 TRM cells in Tmem119 D^b^ cKO mice following VP2_121-130_ injection was accompanied by attenuated BBB disruption visible by MRI. This finding implies microglia antigen presentation enhances the CD8 TRM cell response and overall immune cell infiltration, resulting in increased BBB permeability^3^. The fact that BBB permeability was not attenuated in Tmem119 D^b^ cKO without FTY720 treatment suggests that brain residing CD8 TRM cells can induce BBB regulation more efficiently when additional infiltrating cell types are not present in brain parenchyma. This form of regulation will be investigated in future studies. Nevertheless, our results are highly relevant to our understanding of CD8 TRM function. Brain CD8 TRM cells have been reported to be present in mouse models and human samples and may have a pivotal role in the development and progression of neurological diseases ^4, 5, 7, 59^. Our findings demonstrate an important role for microglia class I restricted antigen presentation in guiding these memory CD8 T cell populations.

The ability to enhance CD8 T cell responses through MHC class I-restricted antigen presentation through discrete cell types may hold potential for therapeutic strategies in diseases where CD8 T cells are known to play a pivotal role in neuropathology. Such diseases include Cerebral Malaria, Multiple Sclerosis, Alzheimer’s Disease and Narcolepsy ^34^. AAV-based therapies targeting discrete MHC class I molecules on microglia could potentially control disease progression and severity. Although more studies are needed to confirm the dynamics of these interactions in other disease models, our results put forward microglia as a key antigen presenting cell in the context of MHC class I during acute and memory phases of infectious disease, regulating CD8 T cell proliferation and BBB integrity.

## Funding

The authors received funding for this work through NIH Grants R01 NS103212, R01 NS122174, NIH training grant T32 AI170478 and the Center for MS and Autoimmune Neurology at Mayo Clinic through the Eugene and Marcia Applebaum Fellowship.

## Acknowledgments

We would like to thank the NIH tetramer core facility for providing tetramers that were used in this work. Biorender.com was used to create the experimental design figures and schematics.

## Notes

### Competing Interest Statement

The authors have declared no competing interest.

