## Supplemental Figures 1, 2 and 3 for "TMEM119+ microglia MHC class I restricted antigen presentation impacts CD8 T cell memory, effector status, and blood-brain barrier disruption during neurotropic virus infection"

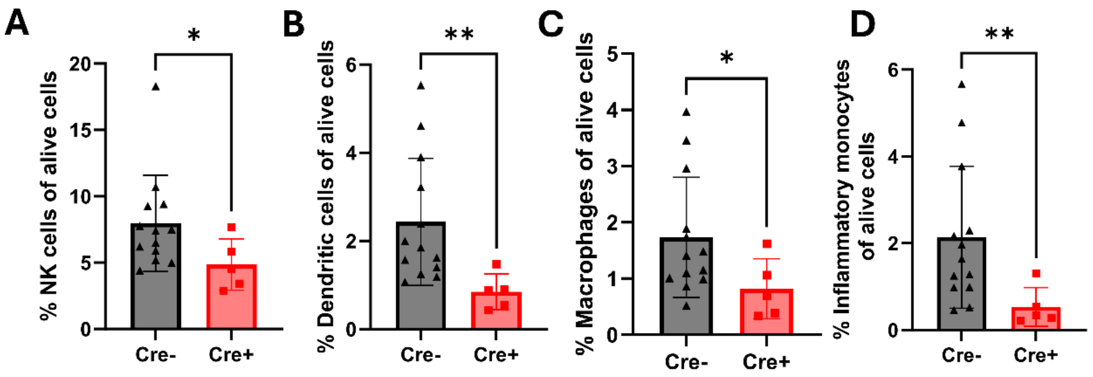


**Figure S1. Reduced frequencies of several immune populations are observed in the Tmem119 K$^b$ cKO 7 days post TMEV-OVA infection.** (A) Bar chart show reduced frequencies of NK cells (TCRβ- CD19- NK1.1+), (B) Dendritic cells (TCRβ- CD19- NK1.1-, CD11c+MHC II+), (C) Macrophages (TCRβ- CD19- NK1.1- F480+) and (D) Inflammatory monocytes (TCRβ- CD19- NK1.1- CD11b+ Ly6C+) in the Cre+ group. Designation of symbols is as follows: ns or not shown for $p > 0.05$, * for $p \leq 0.05$, ** for $p \leq 0.01$, *** for $p \leq 0.001$, **** for $p \leq 0.0001$. Data presented as mean +/- SD.

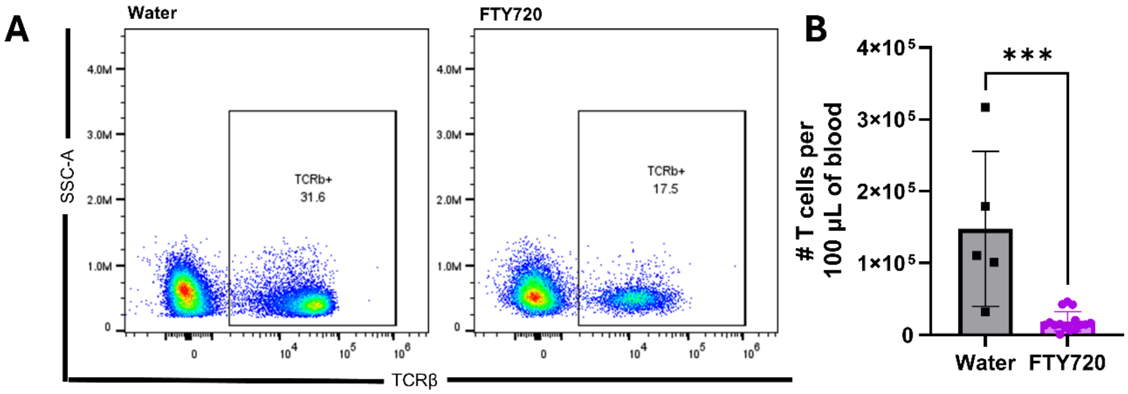


**Figure S2. FTY720 treatment reduces T cell numbers in the blood at 7 days post TMEV infection.** (A) Representative flow cytometry plots show differences between a water control mouse and a FTY720 injected mouse. (B) Bar charts show reduced numbers of T cells per 100 µL of blood in the FTY720 group. Designation of symbols is as follows: ns or not shown for $p > 0.05$, * for $p \leq 0.05$, ** for $p \leq 0.01$, *** for $p \leq 0.001$, **** for $p \leq 0.0001$. Data presented as mean +/- SD.

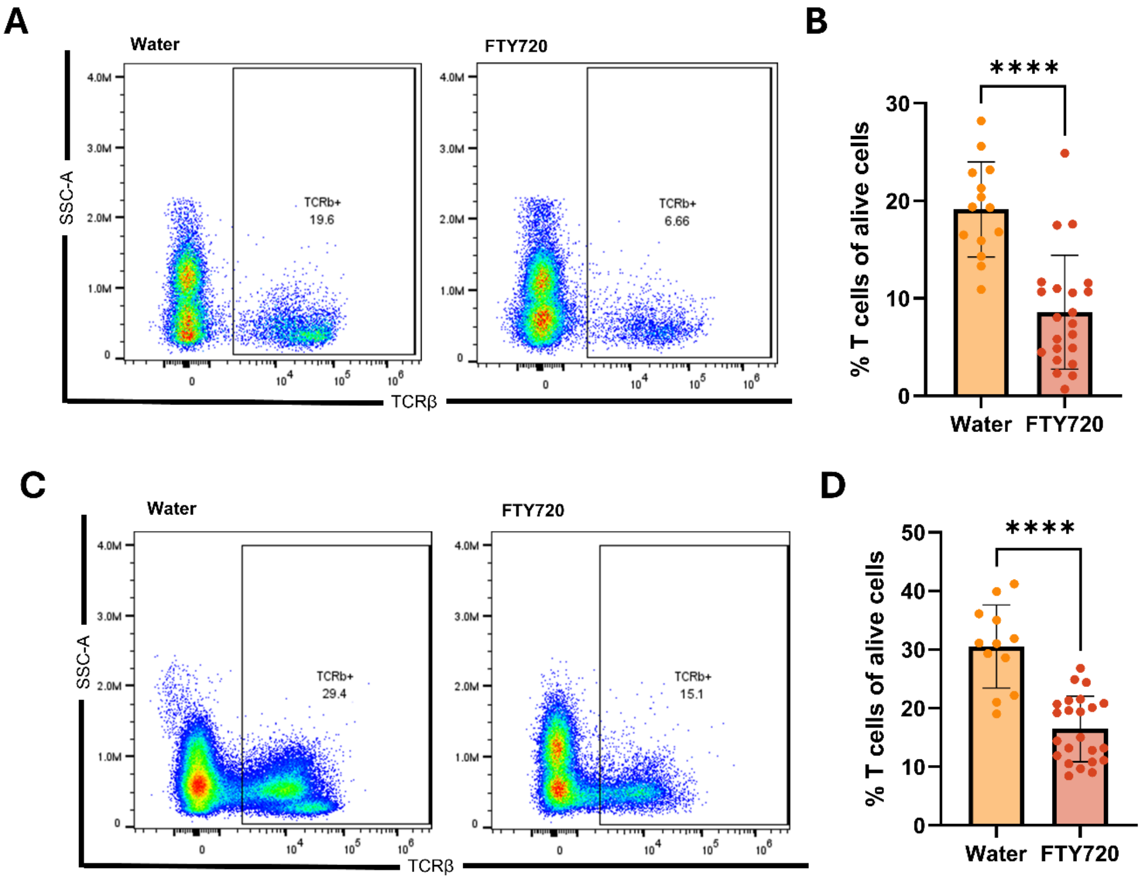


**Figure S3. T cell frequencies are reduced in the blood after 5 days of treatment in a memory reactivation experiment.** (A, B) First cohort of mice used to assess immune cell infiltration 48 hours post $VP2_{121\text{-}130}$ injection. (A) Representative flow cytometry plots show differences between a water control mouse and a FTY720 injected mouse. (B) Bar charts show reduced frequencies of T cells in the blood in the FTY720 group. (C, D) Second cohort of mice used for T1 gadolinium enhanced MRIs 48 hours post $VP2_{121\text{-}130}$ injection. (C) Representative flow cytometry plots show differences between a water control mouse and a FTY720 injected mouse. (D) Bar charts show reduced frequencies of T cells in the blood in the FTY720 group. Designation of symbols is as follows: ns or not shown for $p > 0.05$, * for $p \leq 0.05$, ** for $p \leq 0.01$, *** for $p \leq 0.001$, **** for $p \leq 0.0001$. Data presented as mean +/- SD.
